# *TSniffer*: Unbiased *de novo* identification of RNA editing sites and quantification of editing activity in RNA-seq data

**DOI:** 10.1101/2024.11.13.621086

**Authors:** Maike Herrmann, Yvonne Krebs, Francisco M Acosta, Sebastian Parusel, Oliver Siering, Felix G. M. Andres, Biruhalem Taye, Csaba Miskey, Christian K. Pfaller

## Abstract

RNA editing by adenosine deaminases acting on RNA (ADARs) is an evolutionarily conserved posttranscriptional modification essential for organismal development and normal cell function. Three catalytically active ADARs are conserved in mammals: two isoforms of ADAR1 referred to as ADAR1-p150 and ADAR1-p110, as well as ADAR2. All recognize and edit double-stranded RNA (dsRNA) structures but demonstrate target specificity and selectivity that dictate the unique essential biological functions of the three enzymes. The editing activity of ADAR1-p150 suppresses autoimmune responses against self-RNA structures, whereas ADAR1-p110 and ADAR2 have other primary functions. To better understand the mechanism of target selection by ADARs, we developed *TSniffer*, which allows accurate *de novo* identification of edited transcripts and quantification of the extent of editing within each transcript. We found that 17-40% of protein coding transcripts in mice, ferrets, and humans are edited by ADARs. Individual transcripts can harbor hundreds and thousands of editing sites, mostly within inverted retrotransposable elements. For human transcripts, we found differential editing by ADAR1 and ADAR2, aligning with a supportive role for ADAR2, while some targets were dominantly edited by ADAR1. Relying only on RNA-seq data and reference genome, *TSniffer* represents a novel tool to decipher the role of ADAR editing in different physiological states including disease models. Its unbiased approach is suitable for any organism.

## Introduction

RNA editing by adenosine deaminases acting on RNA (ADARs) is one of the most widespread and evolutionarily conserved forms of post-transcriptional RNA modifications ^1, 2^. There are three catalytically active ADARs in mammals: the two isoforms ADAR1-p150 and ADAR1-p110 are expressed from the *ADAR* gene ^3, 4^, and ADAR2 is expressed from the *ADARB1* gene ^5^. All three enzymes bind to double-stranded RNA (dsRNA) substrates and convert adenosine (A) residues in these substrates to inosine (I) by hydrolytic deamination, a process referred to as A-to-I editing ^6, 7^. Inosine in RNA exhibits base-pairing properties of guanosine (G), and thus, A-to-I editing can alter the coding capacity of RNA as well as dsRNA secondary structures ^8^. ADARs recognize dsRNA structures formed by host or viral transcripts ^9^ and each enzyme exhibits a target selectivity and specificity resulting in unique essential functions of their editing activity ^10–13^.

ADAR editing has numerous implications in health and disease ^14–17^. The essential function of ADAR2 is the site-specific editing of the glutamate ionotropic receptor AMPA type subunit 2 (*GRIA2*) converting the CAG codon for glutamine 607 into a CIG codon for arginine (Q/R site), which is required for receptor function ^18–20^. Although ADAR2 possesses a large number of editing sites across the transcriptome ^21^, disease-causing ADAR2 deficiency in mice surprisingly can be rescued by the single A-to-G mutation at the Q/R site ^22^. In humans, mutations in ADAR2 are associated with neurodevelopmental disorders and seizures ^23, 24^.

In contrast, the biological function of ADAR1, and specifically of its interferon-inducible isoform ADAR1-p150 ^4, 8^, is destabilization of otherwise immunostimulatory cell-derived dsRNAs ^7, 9, 25^. ADAR1 deficiency is lethal in mice and can be rescued by deletion of either the innate immune dsRNA sensor melanoma differentiation-associated gene 5 (MDA-5) ^26–29^, or its downstream adapter mitochondrial antiviral-signaling protein (MAVS) ^30, 31^, or protein kinase R (PKR) ^32^. In humans, mutations altering the activity of ADAR1 are associated with the type-I interferonopathy Aicardi-Goutières-Syndrome (AGS) ^33^. ADAR1 deficiency leads to aberrant activation of dsRNA innate immune sensors including MDA-5, PKR, ribonuclease L, and Z-DNA binding protein 1 (ZBP1), causing unregulated type-I interferon (IFN) responses ^32, 34–40^. Aberrant editing by ADAR1 has implications in cancer development and cancer therapies ^41–44^. ADAR editing mostly occurs in inverted retrotransposable elements such as small interspersed nuclear elements (SINEs), including Alu repeats ^45, 46^.

Recent advancements in computational approaches have revealed insights into the dynamics of RNA editing across different tissues and cell types, as well as to the contributions of individual ADARs to the cellular editome ^12, 21, 47^. However, to more fully understand the implications of different ADARs in disease prevention and development, a detailed categorization of ADAR target transcripts and quantification of the RNA editing levels in individual transcripts is needed ^48^. For this purpose, we have developed and describe herein *TSniffer (Transition Sniffer)*, a bioinformatics tool that allows unbiased *de novo* identification of RNA editing sites in RNA-sequencing (RNA-seq) datasets based on the hyperediting activity of ADARs. In contrast to existing algorithms, *TSniffer* does not depend on availability of RNA editing databases, such as REDIportal ^49^, or the comparison of differential RNA editing in two samples ^50^. Using RNA-seq datasets from wild type (WT) and ADAR-deficient mouse and human samples, we provide a quantitative catalogue of ADAR target transcripts. Differential analyses of transcriptomes from ADAR1-deficient and ADAR1/2-deficient cells provide evidence for target specificity and selectivity of the different enzymes. We also identify RNA editing targets in ferrets providing further evidence for the evolutionary conservation of RNA editing mechanisms across mammalian species.

## Results

### *TSniffer* accurately identifies ADAR hyperediting regions in RNA-seq data

To identify the ADAR-target transcripts, and to quantify the level of editing within these transcripts, we have developed *TSniffer*. The *TSniffer* tool relies on the hyperediting activity of ADARs in dsRNA structures, which can be observed as A-to-G (AG) or T(U)-to-C (TC) transitions (Ts) in RNA-sequencing datasets, dependent upon the transcript-encoding DNA strand. *TSniffer* uses BAM alignments of RNA-seq data as input (**Fig. 1a**) and first generates read count tables (**Fig. 1b**). Using a sliding window approach, *TSniffer* applies Fisher’s exact tests to determine windows with significant enrichment of query transitions (AG, CT, GA, TC) over other occurring mutations (**Fig. 1c**), and merges overlapping significant windows into transition regions (TsRegions; **Fig. 1d**). The GFF-formatted output summarizes the results for each TsRegion including type of transition (TsType), relative transition frequency (RTF), and the number of edited sites (TsSites; **Fig. 1e**). *TSniffer* can be used in two modes: *TSniffer deNovo* analysis; and, *TSniffer Regio* analysis. *TSniffer deNovo* analysis allows initial identification of TsRegions and only requires BAM alignment and the FASTA reference file (**Fig. 1f**); *TSniffer Regio* analysis determines editing activity in a pre-defined set of genomic regions provided as a GFF-formatted file (**Fig. 1f**). This permits quantification and comparison of editing between different samples.

**Fig. 1:**
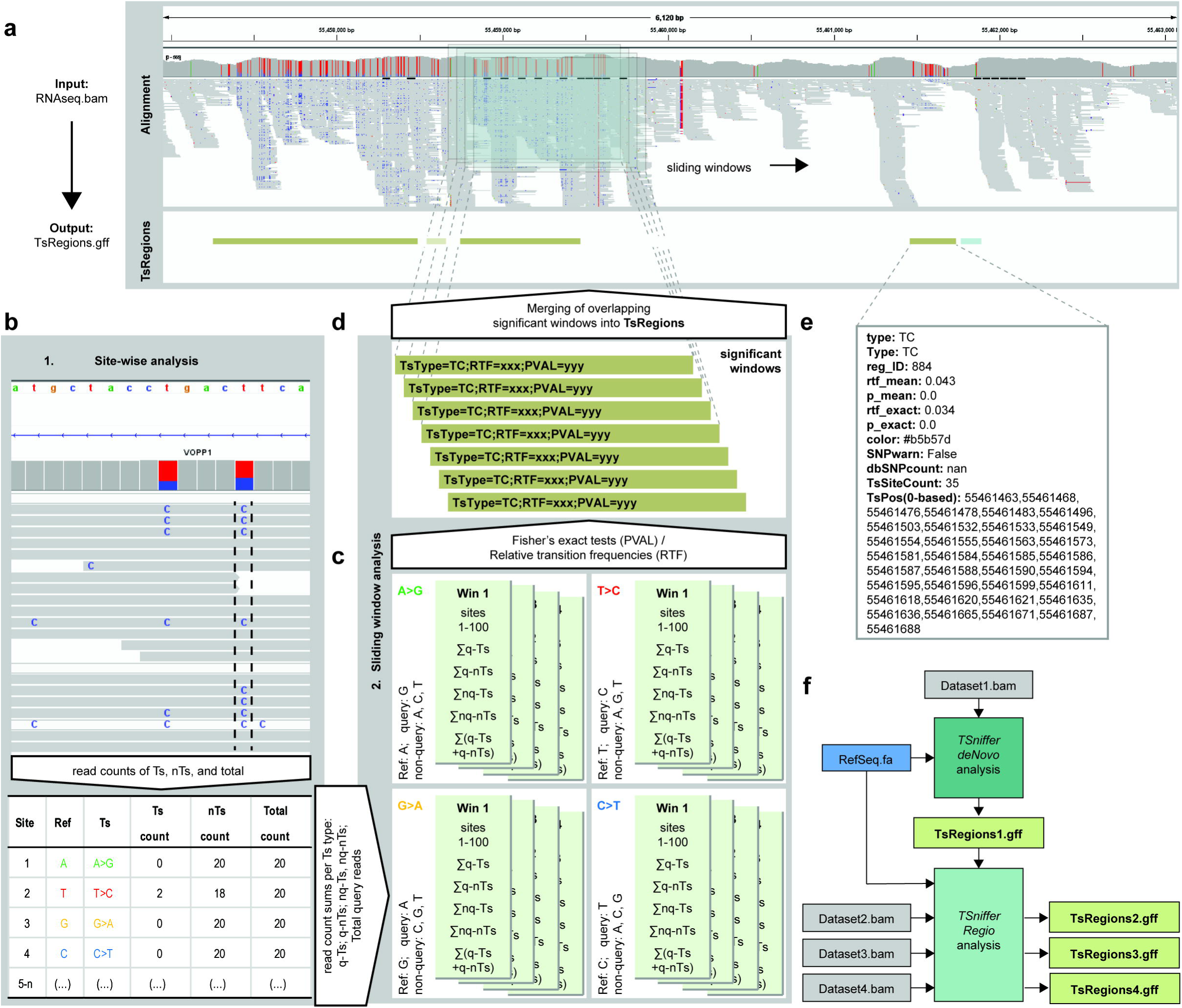
*TSniffer* concept and basic operation modes. (**a**) Screenshot of RNA-sequencing data shown in the integrated genome viewer (IGV). Alignment shows coverage plot (allele frequency threshold is set to 0.05) and aligned reads from imput BAM file. Below is the loaded output GFF file from *TSniffer* analysis. Detected TsRegions are shown as color-coded bars (gold: TC; teal: AG). (**b**) *TSniffer* first performs a site-wise analysis generating read count tables from the input BAM alignment summarizing target transition (Ts) and non-transition (nTs) counts for each position. (**c**) For each transition (Ts) type (AG, CT, GA, TC), *TSniffer* performs a sliding window analysis calculating significant occurrence of Ts types using Fisher’s exact test of query transitions (q-Ts) against query non-transitions (q-nTs), non-query transitions (nq-Ts), and non-query non-transitions (nq-nTs). For details see Methods section. (**d**) Overlapping significant windows are merged into TsRegions. RTF values, *p*-values, and TsSite counts are re-calculated for each window. (**e**) Summary information of a TsRegion. RTF_mean and *p*_mean are derived from the individual overlapping windows, whereas RTF_exact and *p*_exact are re-calculated for the final window. TsPos indicates the exact nucleotide positions of identified TsSites. (**f**) Simple workflow of *TSniffer deNovo* and *TSniffer Regio* analyses.

To establish a workflow to define the transcripts affected by ADAR editing in mice, we used publicly available RNA-seq datasets from WT and ADAR1- and ADAR2-deficient (DKO) mouse brains, which have been shown to possess a high number of ADAR editing sites ^51, 52^. *TSniffer deNovo* identified approximately 12,000 TsRegions in each WT dataset, and about 5,000 TsRegions in each DKO dataset (**Extended Data Fig. 1a**), harboring approximately 70,000 / 20,000 TsSites, respectively (**Extended Data Fig. 1b**). Target TsRegions and TsSites (AG, TC) were 7.5- and 11-fold overrepresented over non-target TsRegions and TsSites (CT, GA) in WT samples, while not enriched in DKO samples (**Extended Data Fig. 1c, d**).

Discarding TsRegions with less than 5 TsSites improved the target to non-target ratio (**Extended Data Fig. 1e, f**). The number of TsRegions with at least 5 TsSites was about 40% of the total number of TsRegions in WT samples, but less than 20% in DKO samples (**Extended Data Fig. 1a**). Nevertheless, this subset of TsRegions maintained about 2/3 of all TsSites (**Extended Data Fig. 1b**), indicating that this constraint is suitable to accurately filter out sequencing/alignment artifacts from ADAR hyperediting regions.

We next determined how sequencing depth affected *TSniffer* performance. For this, we merged the three replicate BAM files of WT and DKO mouse brains into WT_merged and DKO_merged, respectively, and performed *TSniffer deNovo* analysis (**Extended Data Fig. 2a**). As expected, the number of identified TsRegions increased 2-fold (**Fig. 2a**) and of TsSites by a factor of 3 (**Fig. 2b**), while the ratios of target to non-target transitions remained identical (**Extended Data Fig. 2b, c, f, g**). The WT sample exhibited a population of target TsRegions that differed from non-target TsRegions in length, RTF, and number of TsSites (**Extended Data Fig. 2d, h**), while remaining TsRegions detected in the DKO sample showed no difference between target and non-target Ts (**Extended Data Fig. 2e, i**), indicating that these remaining TsRegions in the DKO sample consisted of artifacts. To simplify the annotation and to filter out artificial TsRegions, we first merged TsRegions of the same type that were within 100 nt distance from each other into single TsRegions, and then removed those TsRegions from the WT dataset that had at least 50% overlap with matching TsRegions in the DKO dataset (**Extended Data Fig. 2j**). This filtered set of TsRegions (TsReg_f05) was used for *TSniffer Regio* analysis on the individual datasets, indicating high consistency of detection of TsRegions across the samples (**Extended Data Fig. 2k, l**). However, in each sample about 20-30% of TsRegions were not analyzable due to low coverage (**Extended Data Fig. 2k**). We therefore performed *TSniffer Regio* analysis on the merged WT and DKO datasets, thereby increasing the fraction of analyzable TsRegions to 27,000 (**Fig. 2c, Extended Data Fig. 2m**). To remove additional potential artifacts, we filtered out TsRegions that contained less than 5 TsSites in the WT dataset (**Fig. 2c**). Finally, we calculated a confidence indicator (CI) for each TsRegion based on the reduction of RTF value between WT and DKO sample and filtered for different CI levels (**Fig. 2c**). This additional filtering only mildly affected the number of remaining TsRegions, indicating that our approach was accurate. For downstream analysis, we kept the Mouse_85-5 set of TsRegions with CI ≥ 0.85 and TsSite count ≥ 5, consisting of 13,113 target and 496 non-target TsRegions. The target TsRegions within this dataset exhibited WT-specific enrichment of RTF and TsSite counts relative to TsRegion length compared to non-target TsRegions (**Fig. 2d, e**).

**Fig. 2:**
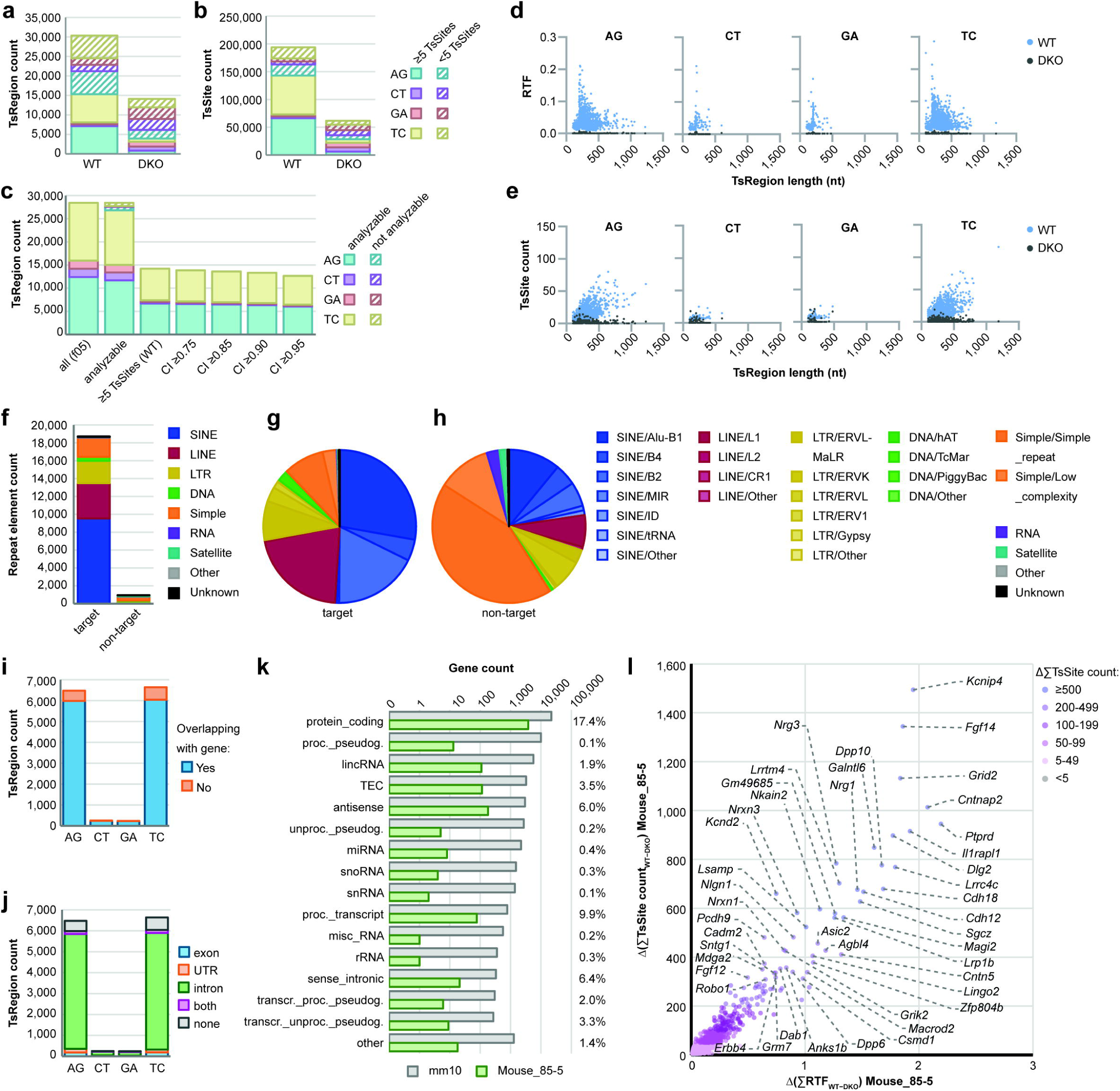
ADAR editing occurs in transcripts of more than 4,300 mouse genes. (**a**) TsRegion count and (**b**) TsSite count of *de novo*-identified TsRegions in merged BAM alignments of wild type (WT) and ADAR1/2-deficient (DKO) mouse brains. (**c**) Filtering of TsRegions by coverage (not analyzable TsRegions had no coverage in at least one dataset), TsSite count (cutoff value: ≥5 in WT), and confidence indicator (CI; cutoff value: ≥0.85 was chosen for downstream analyses and assigned the name Mouse_85-5 set of TsRegions). (**d**) Distribution of RTF values and TsRegion length in the Mouse_85-5 set of TsRegions. WT values are shown in blue, DKO values in dark grey. Each TsType is shown in a separate diagram. (**e**) Distribution of TsSite counts and TsRegion length in the Mouse_85-5 set of TsRegions. (**f**) Number of repeat elements overlapping with TsRegions. Target TsRegions are AG and TC, non-target TsRegions are CT and GA. Colors indicate different repeat families. (**g**) Relative abundance of repeat family subtypes among target TsRegion-containing repeats. (**h**) Relative abundance of repeat family subtypes among non-target TsRegion-containing repeats. (**i**) TsRegion count by intersection with annotated genes. (**j**) Distribution of TsRegions within transcript elements. (**k**) Total number of annotated genes in the mm10 reference genome (grey), total number of genes with target TsRegions (green), and relative frequency of TsRegion-harboring genes, per gene type. (**l**) Quantification of ADAR editing in mouse transcripts. Relative transition frequency (RTF) values and TsSite counts of all Mouse_85-5 TsRegions intersecting with individual genes in WT were summed up and residual values of respective regions in DKO were subtracted. Transcripts are color-coded by total number of calculated TsSites. Most highly edited transcripts are indicated.

### Identification of over 4,000 highly edited mouse brain transcripts by *TSniffer*

Having established a set of mouse TsRegions that was specific for ADAR editing, we next aimed to characterize the genomic features these TsRegions overlapped with, including expressed genes, exon/intron sequences, and integrated repeat elements. For this, we intersected the TsRegion dataset with reference annotation and RepeatMasker files containing the specific genomic features using Bedtools (**Extended Data Fig. 2n**). Over 95% of target TsRegions of the Mouse_85-5 dataset overlapped with annotated repeat elements, while the enrichment of non-target TsRegions within repeat elements was slightly lower (**Extended Data Fig. 3a, b**). The *de novo*-generated set of TsRegions from the WT and DKO samples showed a similar enrichment (**Extended Data Fig. 3c-f**), but only the WT sample had significantly more target TsRegions than non-target TsRegions. We then classified the types of repeat elements overlapping with target or non-target TsRegions. Of over 5 million repeat elements annotated in the mm10 reference genome (**Extended Data Fig. 3g, h**), about 19,000 were found to harbor target TsRegions, and about 1,000 non-target TsRegions from the Mouse_85-5 dataset (**Fig. 2f**). Small interspersed nuclear elements (SINEs), specifically SINE/Alu-B1 and SINE/B2 elements, were enriched for target TsRegions (**Fig. 2g**), whereas non-target TsRegions were mostly found in simple repeat elements (**Fig. 2h**). Similar results were obtained for the WT *de novo* set of TsRegions, whereas in the DKO *de novo* analysis, simple repeat elements were enriched for both target and non-target TsRegions (**Extended Data Fig. 3i-n**). Other notable repeat elements in which ADAR editing occurred were long interspersed nuclear elements L1 (LINE/L1) and long terminal repeats from murine endogenous retroviruses type L-MaLR (LTR/ERVL-MaLR). A more detailed subtype analysis revealed that on average, 0.1-1% of repeat elements from most subtypes were affected by ADAR editing (**Extended Data Fig. 3o**). These data extend previous studies on the preferred editing of SINEs, LINEs, and LTRs by ADARs.

We next determined the transcripts affected by ADAR editing. For this, we identified the annotated genes intersecting with the different sets of TsRegions. About 92% of TsRegions in the Mouse_85-5 dataset were within annotated genes (**Fig. 2i**). Of 55,401 annotated genes in mm10, we identified 2,017 genes to harbor one TsRegion and 2,360 genes to harbor multiple TsRegions, up to 82 (**Extended Data Fig. 4a**). A similar distribution of TsRegions from the WT *de novo* dataset was found (**Extended Data Fig. 4b**), while fewer genes harbored TsRegions from the DKO *de novo* dataset (**Extended Data Fig. 4c**). Consistently between the different datasets, most TsRegions were found in intronic sequences, while untranslated regions (UTRs) and exons were less frequently edited (**Fig. 2j** and **Extended Data Fig. 4d**). 17.4% of all protein coding genes were affected by ADAR editing, while other gene types such as long non-coding RNAs and antisense genes were less frequently affected (**Fig. 2k**). To quantify the overall extent of editing for each transcript, we summed up the RTF and TsSite count values for all TsRegions within each gene from the WT_merged dataset, and subtracted the values determined in the DKO_merged dataset. This allowed identification of highly differentially edited transcripts between WT and DKO samples (**Fig. 2l** and **Extended Data Fig 4e**). Importantly, the Mouse_85-5 subset of TsRegions revealed the same transcripts as the direct analysis of the *de novo* identified TsRegion datasets. While the calculated Δ(∑RTF) values were about 50% reduced using the Mouse_85-5 dataset compared to *de novo* (**Extended Data Fig. 4f**), the numbers of identified TsSites per transcript correlated very well with both approaches (**Extended Data Fig. 4g**). These data indicate that the Mouse_85-5 dataset is highly representative of the entire mouse editome and thus allows to accurately determine the editing levels within individual transcripts. To compare *TSniffer‘s* performance with other methods, we performed a meta-analysis of the edited transcripts identified here with those identified by Tan et al. ^21^. While *TSniffer* detected 187 of 351 previously identified transcripts, it also discovered 4,184 new transcripts to be ADAR editing targets, thereby increasing the landscape of editing in mice by more than 10-fold (**Extended Data Fig. 4h**). To summarize thus far, we have developed *TSniffer*, which detects ADAR hyperediting regions in transcriptomic datasets and allows quntifiction of RNA editing activity. We present various downstream analyses pipelines and show their accuracy by comparing transcriptomes of WT and ADAR-deficient mouse brains. Finally, our approach identified a set of over 4,000 highly edited transcripts in mouse brains.

### ADAR2 has a supporting role in hyperediting of a majority of human ADAR target transcripts

We recently reported identification of transcripts edited by ADAR1 isoforms in human cell lines ^37^. In the analysis of ADAR1-deficient HeLa cells (HeLa 1KO), we found a large number of remaining hyperediting clusters, indicating an involvement of ADAR2. To test whether ADAR2 was responsible for these editing events, we generated ADAR1- and ADAR-double knockout HeLa cells (HeLa DKO; **Extended Data Fig. 5a**) and performed *TSniffer* analysis on HeLa WT, 1KO, and DKO RNA-seq datasets. We employed the same strategy as in the analysis of mouse datasets (**Extended Data Fig. 2a, j, m, n**). *TSniffer deNovo* analysis of the merged BAM alignments of HeLa WT, 1KO, and DKO transcriptomes, respectively, exhibited a moderate reduction of target TsRegion and TsSite counts in HeLa 1KO cells compared to HeLa WT, and a strong reduction in HeLa DKO cells (**Extended Data Fig. 5b-d**), indicating an involvement of ADAR2 to hyperediting of a majority of TsRegions. After removal of TsRegions that were found in the HeLa DKO dataset from the HeLa WT dataset, we identified about 120,000 TsRegions in the resulting HeLa_TsReg_f05 dataset, of which more than 80% were target TsRegions.

The overall coverage in the *TSniffer Regio* analysis of the merged HeLa WT, 1KO, and DKO datasets was much improved compared to the individual samples (**Fig. 3a, Extended Data Fig. 5e**). After removal of TsRegions with low TsSite counts and CI, we defined a set of 49,231 target TsRegions and 876 non-target TsRegions, defined as HeLa_85-5 set of TsRegions (**Fig. 3b**). The TsRegion length varied for target TsRegions from 100 nt to 2,200 nt but remained below 500 nt for non-target TsRegions (**Fig. 3c, d**). The median RTF values (**Fig. 3c**) and TsSite counts (**Fig. 3d**) decreased by 56% and 33% between HeLa WT and 1KO, respectively. The DKO dataset exhibited no ADAR editing activity (100% reduction of median RTF values and TsSite counts compared to WT cells). About 75% of target TsRegions were found within SINE/Alus (**Fig. 3e, f**), whereas non-target TsRegions were less frequently associated with SINE/Alus (**Fig. 3e, g**).

**Fig. 3:**
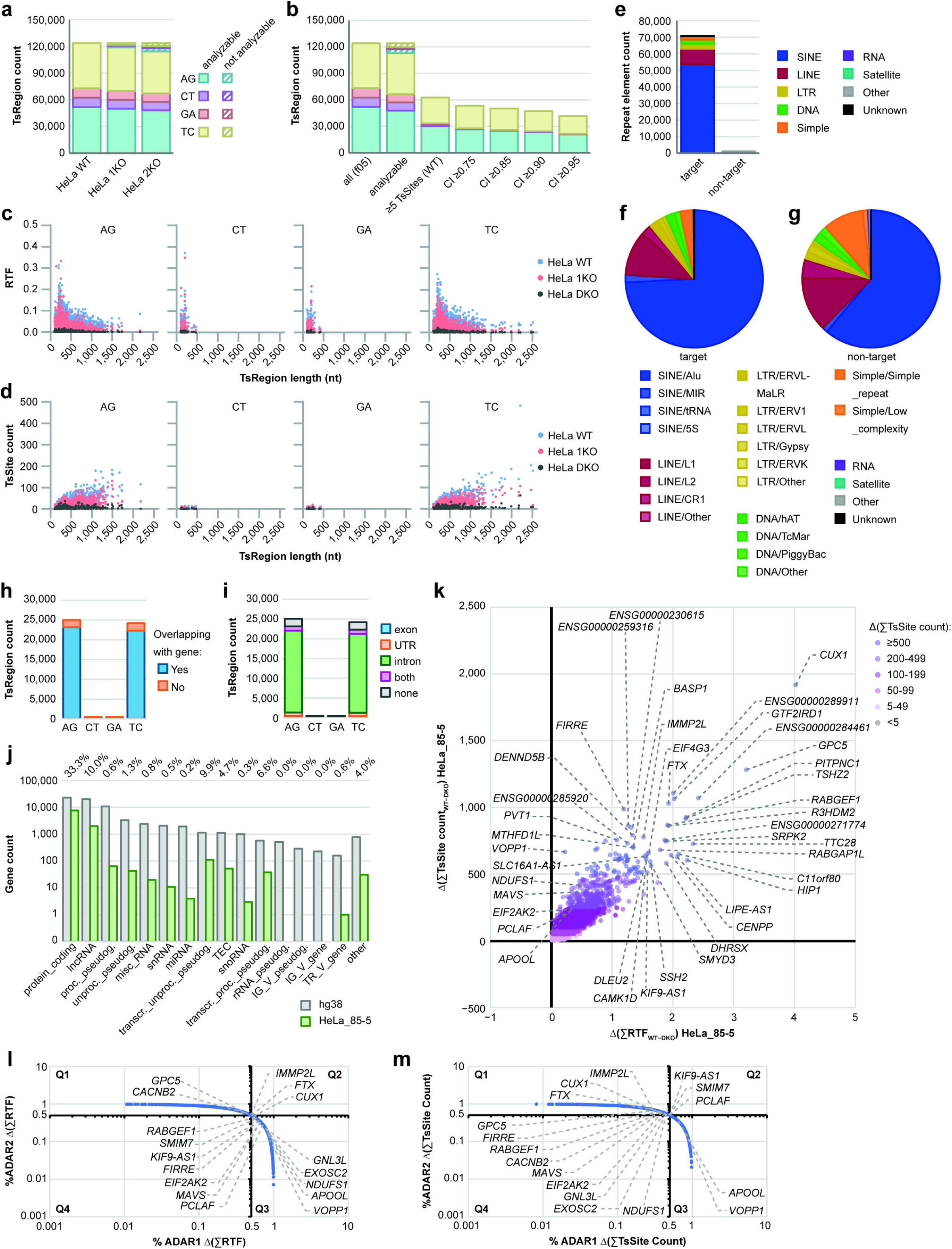
Differential editing of ADAR1 and ADAR2 in human transcipts. (**a**) TsRegion count of WT-specific TsRegions in wild type (WT), ADAR1-deficient (1KO) and ADAR1/2-deficient (DKO) HeLa cells. Not analyzable TsRegions had no coverage in the respective dataset. (**b**) Filtering of TsRegions by coverage (not analyzable TsRegions had no coverage in at least one dataset), TsSite count (cutoff value: ≥5 in WT), and confidence indicator (CI; cutoff value: ≥0.85 was chosen for downstream analyses). (**c**) Distribution of RTF values and TsRegion length in the HeLa_85-5 set of TsRegions. WT values are shown in blue, 1KO values in red, and DKO values in dark grey. Each TsType is shown in a separate diagram. (**d**) Distribution of TsSite counts and TsRegion length in the HeLa_85-5 set of TsRegions. (**e**) Number of repeat elements overlapping with TsRegions. Target TsRegions are AG and TC, non-target TsRegions are CT and GA. Colors indicate different repeat families. (**f**) Relative abundance of repeat family subtypes among target TsRegion-containing repeats. (**g**) Relative abundance of repeat family subtypes among non-target TsRegion-containing repeats. (**h**) TsRegion count by intersection with annotated genes. (**i**) Distribution of TsRegions within transcript elements. (**j**) Total number of annotated genes in the hg38 reference genome (grey), total number of genes with target TsRegions (green), and relative frequency of TsRegion-harboring genes, per gene type. (**k**) Quantification of ADAR editing in human transcripts. Relative transition frequency (RTF) values and TsSite counts of all HeLa_85-5 TsRegions intersecting with individual genes in WT were summed up and residual values of respective regions in DKO were subtracted. Transcripts are color-coded by total number of calculated TsSites. Most highly edited transcripts are indicated. (**l**) Relative contribution of ADAR1 and ADAR2 enzymes to calculated RTF values per transcript. (**m**) Relative contribution of ADAR1 and ADAR2 enzymes to calculated total TsSite counts per transcript.

Compared to the total occurrence of different repeat elements in the human reference genome hg38 (**Extended Data Fig. 6a, b**), this indicated a strong association of ADAR editing with SINE/Alus and confirms previous studies ^53^. However, significant editing was also found in some LINE, LTR, and DNA transposon repeats (**Extended Data Fig. 6c**). Over 90% of TsRegions were found in annotated genes (**Fig. 3h**), and similar to our results of the mouse transcriptome analysis, most of editing regions were found in introns (**Fig. 3i**). After assigning each TsRegion to their overlapping gene products, we found that one third of all protein coding genes (7,615) and one tenth of long non-coding RNAs (lncRNA; 1,999) were found to harbor ADAR editing TsRegions (**Fig. 3j**). Transcripts of other gene types were less frequently targeted by ADARs. The number of total TsRegions per gene ranged from one to 135, with 34 genes possessing 50 or more TsRegions (**Extended Data Fig. 7a**).

The calculated Δ(∑RTF) and Δ(∑TsSite count) scores of the HeLa_85-5 TsRegion set revealed the most highly differentially edited transcripts between HeLa WT and DKO cells (**Fig. 3k**) Among these, *CUX1* was found to harbor over 1,900 TsSites within 222 TsRegions. *CUX1* is a large gene with a total transcript size of 468 kb (3 kb exons and 465 kb introns). A total of 1,096 repeat elements are integrated in introns, the majority of which are Alu repeats (525). 117 of these Alu repeats (33.7%) were found to harbor TsRegions. Previously described candidates, such as *VOPP1*, *NDUFS1*, *APOOL*, *FIRRE*, and *PCLAF* ^37^, were found to harbor between 100 and 1,000 TsSites. The majority of these TsSites are located in 3’UTRs of these transcripts. The results obtained with the HeLa_85-5 dataset correlated well with the alternative differential analysis of *de novo*-identified TsRegions in HeLa WT and DKO samples, both on the level of RTF (**Extended Data Fig. 7b**) and TsSite count (**Extended Data Fig. 7c**).

We next estimated the relative contribution of ADAR1 and ADAR2 editing to the cumulative editing in each transcript by differential analysis of WT versus 1KO (ADAR1-effect, **Extended Data Fig. 7d**), and 1KO versus DKO (ADAR2-effect, **Extended Data Fig. 7e**). This revealed that ADAR1 and ADAR2 enzymes contributed to editing of the majority of transcripts at various degrees, and knockout of either enzyme partially reduced RTF values (**Fig. 3l**) and TsSite counts (**Fig. 3m**). However, some genes were dominantly edited by ADAR1 (e.g. *VOPP1*, *APOOL*, *NDUFS1*, *EXOSC2*, *GNL3L*, *PCLAF*, *MAVS*, *EIF2AK2*), whereas a different subset was dominantly edited by ADAR2 (*CACNB2*, *GPC5*, *IMMP2L*, *FTX*, *CUX1*). We found a stronger contribution of ADAR1 to editing of genes with large numbers of TsSites (**Extended Data Fig. 7f**), while the contribution of ADAR2 was more apparent in genes with overall lower numbers of TsSites (**Extended Data Fig. 7g, h**). Importantly, transcripts most highly edited by ADAR2 still exhibited a significant contribution of ADAR1, suggesting that ADAR2-exclusive TsRegions are rare. In contrast, we found transcripts with nearly exclusive contribution of ADAR1 to the overall editing. In summary, we have successfully applied *TSniffer* to quantify differential editing of ADAR1 and ADAR2 enzymes in human cell lines with altered expression of ADARs. Our results indicate that ADAR1 uniquely targets a subset of highly edited transcripts whereas ADAR2 possesses a supportive role in editing of most of highly edited transcripts.

### ADAR TsRegions are conserved between cell lines and primary cells

We next asked whether the ADAR hyperediting events we detected in an immortalized human cell line were biologically relevant for primary human tissue. For this, we selected three brain RNA-sequencing datasets from the GTEx database with high sequencing depth ^54^. We chose brain tissue for three reasons: first, brain tissue was reported to have high ADAR editing activity ^21^; second, direct comparison to our analysis of mouse brain tissue (**Fig. 2**) could allow identification of evolutionary conserved target transcripts; and third, expression of transcripts not expressed in HeLa cells would allow expanding the spectrum of human target transcripts.

*TSniffer deNovo* detected between 30,000 and 70,000 TsRegions with 250,000-800,000 TsSites in each dataset (**Extended Data Fig. 8a, b**). The sensitivity of detection was strongly increased by combining the three datasets, resulting in a set of about 70,000 TsRegions with at least 5 TsSites, defined as GTEx_Brain_m100-5 set of TsRegions (**Fig. 4a)** and combining over 1.2 million TsSites highly specific for target Ts (**Fig. 4b**). Consistent with the results from the HeLa cell analysis, target TsRegions were found to overlap with SINE/Alu and LINE/L1 repeats (**Extended Data Fig. 8c, d**), whereas non-target TsRegions did not (**Extended Data Fig. 8c, e**). 85% of target TsRegions overlapped with annotated genes (**Extended Data Fig. 8f**), and the majority of TsRegions were found within introns (**Extended Data Fig. 8g**). The total number of protein coding genes and lncRNAs with target TsRegions was higher than in HeLa cells (**Extended Data Fig. 8h**). Together, these data indicate higher editing activity in primary human brain tissue than in HeLa cells, which may be the result of higher heterogeneity of cell types in brain samples in comparison to a clonal cell line.

**Fig. 4:**
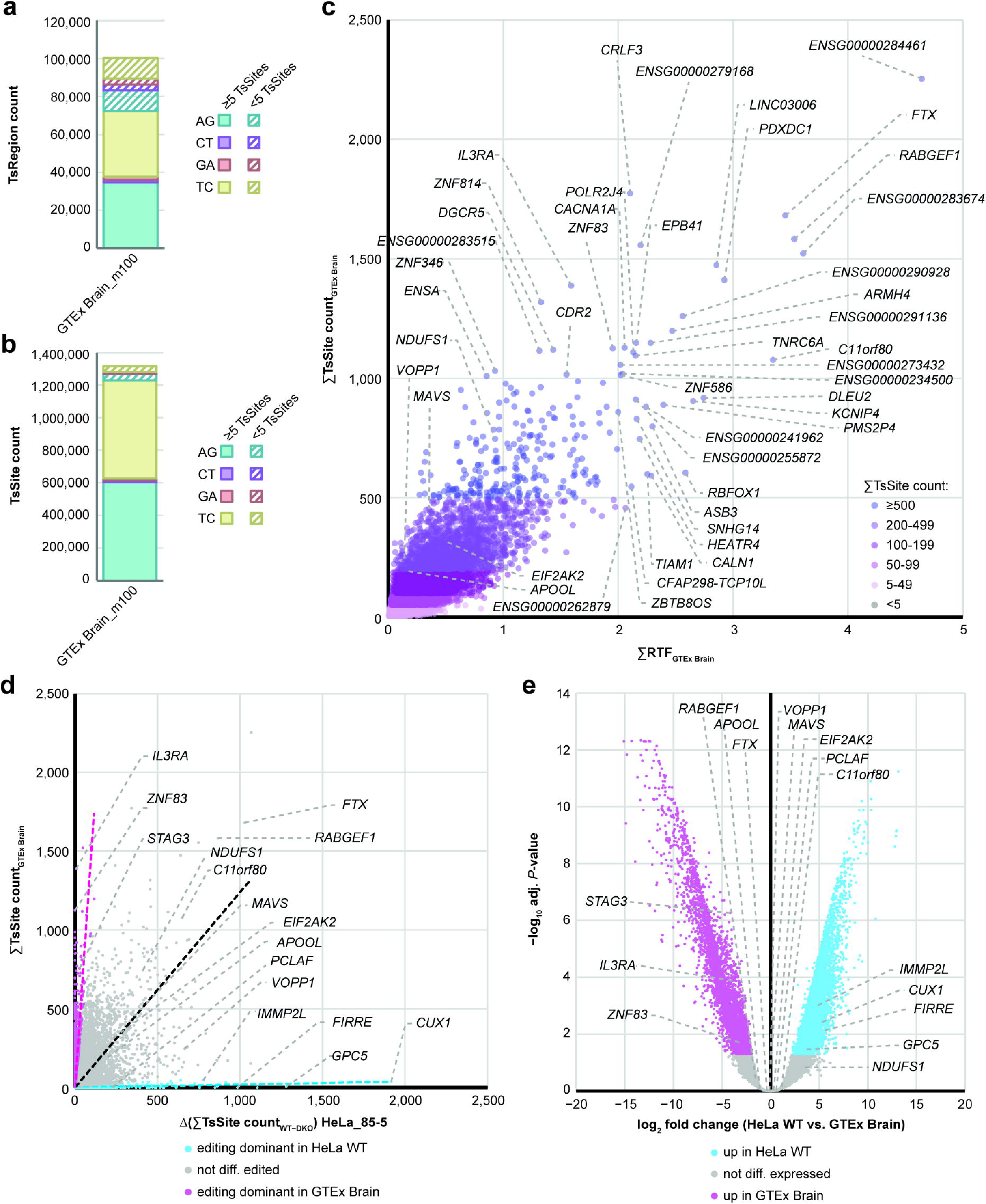
ADAR editing in cell lines is consistent with editing in primary tissue. (**a**) TsRegion count and (**b**) TsSite count of *de novo*-identified TsRegions in the transcriptome of the human brain. (**c**) Quantification of ADAR editing in human brain transcripts. Relative transition frequency (RTF) values and TsSite counts of all Brain TsRegions with ≥5 TsSites (Brain_m100-5) intersecting with individual genes were summed up. Transcripts are color-coded by total number of calculated TsSites. Most highly edited transcripts are indicated. (**d**) Correlation of gene-associated total TsSite counts detected in HeLa WT cells using the HeLa_85-5 set (x-axis) with values from human brain samples using the Brain_m100-5 set (y-axis). Color coding if gene was dominantly edited in HeLa WT (blue; TsSite count[HeLa WT] ≥ 9 × TsSite count[Brain]) or if gene was dominantly edited in brain (pink; TsSite count[Brain] ≥ 9 × TsSite count[HeLa WT]. Dashed lines indicate linear regression for the different groups. (**e**) Differential gene expression analysis in HeLa WT and GTEx brain samples. Genes with significantly higher expression in HeLa WT are indicated in blue, genes with significantly higher expression in brain are indicated in pink.

To directly compare the editing targets in HeLa cells and primary brain tissues, we next performed *TSniffer Regio* analysis of the HeLa WT and the combined GTEx brain alignment using the HeLa_85-5 set of TsRegions (**Extended Data Fig. 9a-c**), and the GTEx Brain_m100-5 set of TsRegions (**Extended Data Fig. 9d-f**). Of the 50,000 TsRegions defined in the HeLa_85-5 set, 19,000 returned analyzable data from the GTEx brain alignment (**Extended Data Fig. 9a**).

Comparing the RTF values (**Extended Data Fig. 9b**) and TsSite counts (**Extended Data Fig. 9c**) of these TsRegions, we found a moderate correlation between HeLa WT and brain samples, although RTF values were overall higher in the brain sample. A similar correlation was obtained when the HeLa WT and brain samples were analyzed using the 50,000 shared GTEx Brain_m100 TsRegions (**Extended Data Fig. 9d-f**), indicating that the editing activities in HeLa WT cells and primary brain tissue had large overlaps.

We next identified the cumulative editing scores in brain transcripts and identified the most highly edited genes (**Fig. 4c**). Among these, we identified several that we had also identified in HeLa cells, such as *FTX*, *C11orf80*, *NDUFS1*, *VOPP1*, and *RABGEF1*. In these cases, direct comparison of cumulative RTF values and TsSite counts correlated well between HeLa WT and primary brain samples (**Fig. 4d**, and **Extended Data Fig. 9g**). Others were exclusively edited in brain tissue, for example *IL3RA*, *ZNF83*, *STAG3* (**Fig. 4d**, and **Extended Data Fig. 9g**), while some transcripts edited in HeLa cells were not identified in brain samples, such as *CUX1*, *IMMP2L*, *FIRRE*, *GPC5* (**Fig. 4d**, and **Extended Data Fig. 9g**). Differential gene expression was identified as the reason for the discrepancies in detection of editing for most of the targets (**Fig. 4e**). In summary, our comparison of detection of RNA editing in primary human brain tissue validated the biological relevance of TsRegions detected in immortalized human cell lines. In addition, by setting a cutoff of ≥5 TsSites per TsRegion, we have established a pipeline to determine RNA editing in cell types or tissues lacking a genetic knockout control.

### *TSniffer* exhibits unmet depth of RNA editing analysis while maintaining high accuracy in determining editing regions

To compare the performance of *TSniffer* with other approaches, we performed a meta-analysis of the transcript discovery rates in our HeLa and brain datasets and in four previous studies ^21, 35, 36, 55^ (**Extended Data Fig. 10a-d**). In all cases, *TSniffer* identified most previously described transcripts in either one or both datasets analyzed here, while only a small fraction between 1.8 and 8.4% of the previously identified genes were not detected by *TSniffer*.

Moreover, we performed a side-by-side comparison of the performance of *TSniffer* with *LoDEI*, a recently published algorithm that detects RNA editing sites using a sliding window approach ^50^. This program compares differential RNA editing in two datasets (HeLa WT_merged and HeLa DKO_merged) in windows of defined length. Of 49,231 target TsRegions of the HeLa_85-5 dataset, 35,687 (72.5%) were also identified by *LoDEI* (**Extended Data Fig. 10e**). TsRegion length determined by *TSniffer* correlated well with the total length of windows found by *LoDEI* (**Extended Data Fig. 10f, h**). Due to its 1 nt sliding window approach, *TSniffer’s* TsRegions exhibited a more dynamic length range than the *LoDEI* windows. However, *LoDEI* also identified 49,763 regions that were not included in our HeLa_85-5 dataset (**Extended Data Fig. 10g**). Most *LoDEI*-specific regions were short (mean merged window length was 115 nt) in comparison to those identified by both *LoDEI* and *TSniffer*, or only by *TSniffer* (**Extended Data Fig. 10i**), suggesting that these regions correlated with those discarded in our downstream filtering process due to low number of TsSites (**Fig. 3b**). Unfortunately, *LoDEI* does not report the exact nucleotide positions contributing to the calculated window editing index (wEI). On gene level, of 9,922 edited genes in HeLa cells identified by *TSniffer*, 9,000 were also identified by *LoDEI*, as well as 4,300 additional genes that were only identified by *LoDEI* (**Extended Data Fig. 10j**). Comparison of the *LoDEI* wEI and *TSniffer* RTF on gene level for those genes determined by both algorithms showed high correlation across the entire spectrum of edited transcripts (**Extended Data Fig. 10k**). In summary, *TSniffer* identified more editing sites and edited genes using a sliding window approach than previously reported site-wise analyses ^21, 35^. Importantly, *TSniffer* achieves this depth of analysis without the need for additional annotations ^56, 57^ or matching with editing site databases such as RADAR or REDIportal ^49, 58^. *TSniffer’s* performance compares to other sliding window approaches ^50^, and our downstream analysis pipeline allows for additional filtering of potential artifacts.

### ADAR editing in ferret PBMCs underlies mechanisms conserved across mammals

Ferrets are an important model organism for infectious disease studies, but the available genomic information for ferrets is still less developed than that of mice and humans. Like other mammals, ferrets express three ADARs with high amino acid sequence and protein domain conservation compared to other mammalian ADARs. The *ADAR* gene encoding ADAR1-p150 and ADAR1-p110 is located on contig NW_004569490.1 (81,699-105,573 bp); the *ADARB1* gene is found on contig NW_004569228.1 (7,805,280-7,884,797). We hypothesize that the mechanisms of ADAR editing in the ferret transcriptome are conserved when compared to other mammals. To test whether *TSniffer* can accurately identify ADAR editing in ferret RNA-sequencing datasets, we performed transcriptomics analyses on RNA from primary blood mononuclear cells (PBMCs) isolated from three animals. To maximize the sensitivity of *TSniffer deNovo*, we combined the three alignment files as in previous experiments and identified over 200,000 TsRegions in total, with 89,000 TsRegions harboring at least 5 TsSites (**Fig. 5a**).

**Fig. 5:**
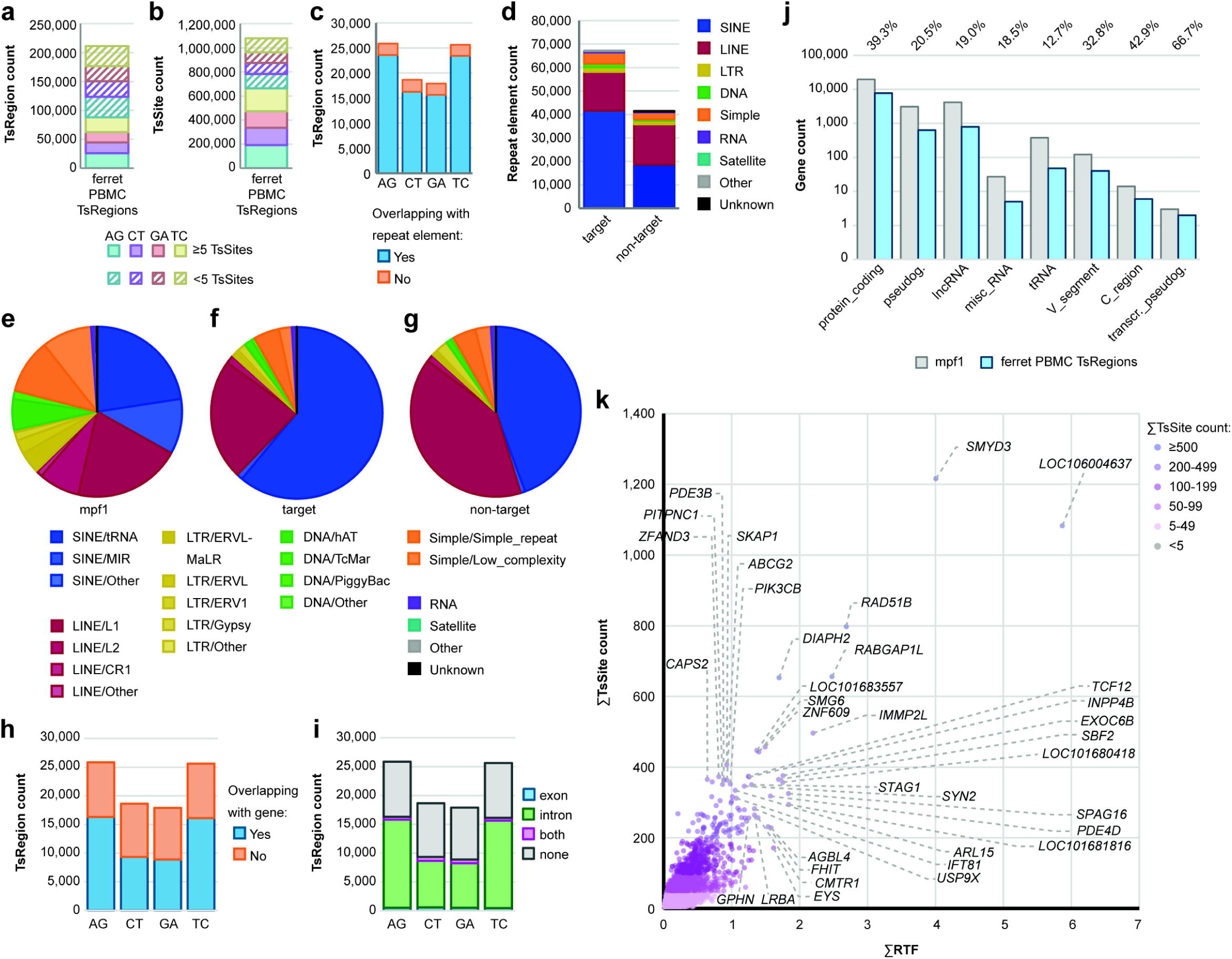
ADAR editing in the ferret transcriptome follows conserved mechanisms. (**a**) TsRegion count and (**b**) TsSite count of *de novo*-identified TsRegions in ferret PBMCs. (**c**) TsRegion count of intersection with annotated repeat elements. (**d**) Number of repeat elements overlapping with TsRegions. Target TsRegions are AG and TC, non-target TsRegions are CT and GA. Colors indicate different repeat families. (**e**) Relative abundance of annotated repeat family subtypes in the ferret reference genome (mpf1). (**f**) Relative abundance of repeat family subtypes among target TsRegion-containing repeats. (**g**) Relative abundance of repeat family subtypes among non-target TsRegion-containing repeats. (**h**) TsRegion count by intersection with annotated genes. (**i**) Distribution of TsRegions within transcript elements. (**j**) Total number of annotated genes in the mpf1 reference genome (grey), total number of genes with target TsRegions (blue), and relative frequency of TsRegion-harboring genes, per gene type. (**k**) Quantification of ADAR editing in ferret transcripts. Relative transition frequency (RTF) values and TsSite counts of all ferret TsRegions with ≥ 5 TsSites intersecting with individual genes were summed up. Transcripts are color-coded by total number of calculated TsSites. Most highly edited transcripts are indicated.

Interestingly, we detected a high number of non-target TsRegions; target TsRegions were only enriched by a factor of 1.4, independent of the filtering by minimum number of TsSites. This indicated a higher mutational background noise likely due to the incomplete assembly of the ferret genome. 62% of target TsSites (about 190,000 each for AG and TC TsSites) and 61% of non-target TsSites (about 143,000 for CT and 137,000 for GA TsSites) were found within TsRegions with at least 5 TsSites (**Fig. 5b**). 92% of target TsRegions (about 47,000) and 87% of non-target TsRegions (about 32,000) were found within annotated repeat elements (**Fig. 5c**).

67,000 individual repeat elements overlapped with target TsRegions, and 41,000 repeat elements overlapped with non-target TsRegions (**Fig. 5d**). In comparison to the overall distribution of repeat element types in the ferret genome (**Fig. 5e**), target TsRegions were mostly associated with SINE/tRNA and LINE/L1 repeats (**Fig. 5f**), aligning with our initial hypothesis. Non-target TsRegions were also enriched in these two types of repeat elements (**Fig. 5g**), but the degree of enrichment was inverse to target TsRegions. The proportion of TsRegions found within annotated genes ranged between 50 and 60% depending on the TsType (**Fig. 5h**) and thus was lower than for mouse (**Fig. 2i**) or human datasets (**Fig. 3h**). However, the distribution of TsRegions between exonic and intronic regions was comparable to the other organisms (**Fig. 5i**). Overall, the fraction of gene types harboring TsRegions exceeded the values found in mice and humans, likely due to the high heterogeneity of cell types present in PBMCs (**Fig. 5j**). The landscape of ADAR-edited genes in ferret PBMCs showed a similar distribution of highly edited transcripts (**Fig. 5k**). Notably, several identified candidates had also been found to be edited in human and mouse samples (e.g. *IMMP2L*, *SMYD3*, *PDE3B*, *ZFAND3*, *STAG1*, *PDE4D*, *SBF2*), indicating evolutionary conservation of editing mechanisms. In summary, we present a quantitatively ranked catalogue of 9,259 annotated ferret genes expressed in PBMCs that harbor TsRegions indicative for ADAR editing.

## Discussion

RNA editing by ADARs is one of the most common post-transcriptional modifications. Editing has widespread biological implications ranging from sequence-dependent effects, such as recoding and alternative splicing of mRNAs and miRNAs ^59, 60^, to sequence-independent effects such as altering the secondary structures of otherwise immunostimulatory dsRNAs ^8, 9, 35, 37, 61, 62^. While the general mechanisms of ADAR editing have been elucidated during the past decades, important questions remain unanswered about the target-selectivity and -specificity of individual ADARs ^48^. Understanding these aspects of target selection will allow us to understand how the three catalytically active mammalian ADARs — ADAR1-p150, ADAR1-p110 and ADAR2 — can mediate individual essential biological functions despite performing the same catalytic reaction on dsRNA substrates. For this purpose, we have developed the tool *TSniffer*, which detects hyperediting clusters, a hallmark of ADAR editing, in next generation RNA-seq datasets. In contrast to other approaches, such as RADAR ^58, 63^, *TSniffer* does not require pre-existing knowledge of editing sites, allowing its use for analysis of editing in any species. Herein, we tested *TSniffer‘s* functionality using RNA-seq datasets from mice, humans, and ferrets.

For both mouse and human data, our analyses confirmed previously identified ADAR editing target transcripts ^28, 35–37^ and increased the number of identified transcripts to over 4,000 in the mouse, and nearly 10,000 in the human transcriptome, while maintaining a rigorous filtering of potential artifacts in the downstream analysis pipeline. Due to its 1 nt sliding window approach, *TSniffer* provided a much more precise identification of ADAR hyperediting positions and clusters than previous analyses that usually relied on individually identified TsSites via tools like JACUSA ^56^ or GIREMI ^57^, or comparison with RNA editing databases such as RADAR ^58^ or REDIportal ^49^. *TSniffer* performed equally well as the recently published sliding window approach *LoDEI* ^50^. However, *TSniffer* offered several advantages: First, our 1 nt sliding window approach offers a more dynamic detection of TsRegions than *LoDEI*, which slices a target sequence into adjacent windows of defined size. Second, *TSniffer* is truly unbiased as it does not require annotation files as input, making it suitable for analysis of RNA-seq datasets from less-well characterized organisms. Third, *TSniffer* allows authentic *de novo* identification and quantification of TsRegions in individual datasets, while *LoDEI* is designed to calculate the differential editing between two datasets. Fourth, *TSniffer* not only provides genomic coordinates of ADAR editing regions and a quantitative measure of the amount of editing, but also includes the position of all sites that contribute to the observed level of editing, which allows implementation of cross-checks with RNA editing databases.

*TSniffer* is perfectly complementary to the *Alu editing index (AEI)* method recently established by Roth et al. ^47^. This method provides a single value for the ADAR editing activity in a sample and allows easy comparison between different tissue types or donors. *TSniffer* calculates exact editing scores for each identified TsRegion. We present a downstream analysis pipeline allowing identification of edited transcripts and transcript-specific quantification of ADAR editing levels. We detected ADAR editing in about one third of protein coding transcripts in all three species analyzed. In addition, ADAR editing was frequently detected in lncRNAs and pseudogenes. Our approach allowed us to rank transcripts by the degree of ADAR editing. A significant portion of genes were edited across the three species, suggesting that ADAR editing targets have co-evolved with their hosts, driven by increased accumulation of retrotransposable elements.

Using mouse and human WT and ADAR1/2-deficient datasets, we show that *TSniffer* accurately detects ADAR-mediated hyperediting clusters. In all three species — mice, humans, and ferrets — editing most frequently occurred in integrated SINEs and LINEs, confirming previous observations ^35–37, 46, 48, 64, 65^. Other repeat elements were also found to be targets of ADAR editing, but to a lesser extent. Importantly, the frequency of editing clusters detected in different species correlated directly with the increased frequencies of SINEs and LINEs in these genomes. In mice, SINE/B1 repeats (the evolutionary analog to human SINE/Alu elements ^45^), SINE/B2 repeats, and LINE/L1 repeats are the dominant targets for ADAR editing. Similarly, SINE/tRNA and LINE/L1 repeats are the dominant targets of ADAR editing in the ferret transcriptome, indicating a long evolutionary conservation of editing targets.

Our study is the first to define the editome of ferrets, determining over 9,000 transcripts harboring ADAR hyperediting clusters. Although *TSniffer* analysis of ferret RNA-seq data generated a higher level of non-target TsRegions indicating a higher mutational noise in the produced BAM alignments, our approach allowed identification of highly edited transcripts. Many of these transcripts were found in samples from all three organisms, supporting the validity of our analyses and suggesting evolutionary conservation of these editing targets.

By comparing the effects of ADAR1-deficiency and ADAR1/2-deficiency in HeLa cells we were able to detect differential effects of ADAR1 and ADAR2 enzymes on editing of specific transcripts. We found that ADAR1 had a dominant role in highly edited transcripts (≥ 150 TsSites per transcript), whereas the relative contribution of ADAR2 increased in transcripts with lower editing rate. Some highly edited transcripts possessed nearly ADAR1-exclusive TsRegions, whereas ADAR2 exhibited a supportive role. These findings align with the essential role of ADAR1 in immunoregulation ^14, 16, 30, 32, 35, 37, 66–69^, while ADAR2 is not required for this function ^22^. However, additional studies will be required to identify transcripts that are exclusively edited by ADAR1-p150 and not by ADAR1-p110, since the former isoform is essential for regulating innate immune responses to dsRNA ^28, 32, 35, 37, 70^. *TSniffer* will assist in identifying the transcripts that are likely associated with autoinflammation ^32, 33, 62^.

In light of the existing dogma that ADAR2 editing is site-specific while ADAR1 editing is promiscuous ^71^, the extent of ADAR2 hyperediting was initially surprising. However, our data align with previous analyses in mouse brains indicating many ADAR2 hyperediting clusters ^52^. Additional analyses revealed that most of the ADAR2-specific editing events occurred in intronic sequences, as exemplified by *CUX1*, *FIRRE*, *IMMP2L*, and *GPC5* transcripts, which were among those with the highest number of ADAR2-specific TsSites. Under normal conditions, introns are efficiently degraded and thus not sensed by dsRNA receptors. Thus, it may not be important whether their internal dsRNA structures were properly edited or not.

However, a recent study showed that introns can be sensed by OAS3/RNase L and PKR and activate an innate immune response ^72^. In addition, spliceosome-targeted cancer therapies were shown to activate dsRNA-mediated innate immune responses through accumulation of mis-spliced mRNAs in the cytoplasm ^73^. Therefore, a role of ADAR2 and ADAR1-p110 in innate immune regulation cannot yet be fully excluded for conditions that would lead to cytoplasmic accumulation of unedited p110/ADAR2 target sequences. Nevertheless, these would be exceptional events, while lack of editing of cytoplasmic dsRNA structures by ADAR1-p150 is the common cause for type-I interferonopathies ^28, 30–33, 35, 36, 74, 75^.

The current state of the art suggests that ADAR1-p150, which is the only ADAR with nucleo-cytoplasmic localization ^74, 76^, is essential for editing of inverted Alu repeats in UTRs of mature mRNAs ^28, 37, 74^. Indeed, many of the human transcripts dominantly edited by ADAR1 harbor TsRegions in their UTRs, for example *APOOL*, *NDUFS1*, *PCLAF*, *MAVS*, or *EIF2AK2*. In some cases, our analysis pipeline incorrectly associated TsRegions to introns rather than extended UTRs spanning outside of the annotated transcript. An example for this is the *VOPP1* transcript. The two highly edited TsRegions (chr7:55,454,193-55,456,705 and chr7:55,457,270-55,459,479) spanning across two inverted LINE/L1 repeats are located about 10 kb downstream of the annotated 3’UTR of regular *VOPP1* transcript variants. However, several predicted *VOPP1* transcript variants possess an intron spanning across the TsRegions and therefore, in our analyses, the *VOPP1*-associated TsRegions were associated with an intron. Combined analyses of splice variant expression levels and the contribution of different ADARs on editing of these variants would allow generation of more precise genome annotations.

In conclusion, *TSniffer* is a novel tool to accurately define ADAR editing regions in RNA-seq datasets and to quantify ADAR editing activity on an individual transcript level. It will aid us in understanding the mechanisms of target selection of different ADARs and identification of immunostimulatory transcripts in autoimmune diseases and cancer. Dysregulated RNA editing has been observed in various cancers and also affects tumor immune escape and the efficacy of cancer immunotherapies ^41^. However, what has been found so far may only be the tip of the iceberg. Furthermore, *TSniffer* is an excellent tool to screen for off-target activities of novel base editors developed for gene therapy ^77^. The dimensions of ADAR editing activity in mammalian transcriptomes suggest that altered editing activity may be involved in many diseases beyond cancer and type-I interferonopathies.

## Materials and Methods

### Cell lines

HeLa cells and derivatives were cultivated in high glucose D-MEM (Sigma-Aldrich, #D6546) supplemented with 10% FBS (Thermo Fisher Gibco, #10500064), 200 mM L-glutamine (Sigma-Aldrich, #G7513), and 10,000 U/l Penicillin, 10 mg/l Streptomycin (Thermo Fisher Gibco, #15140122). Cells were grown in a humified incubator with 5% CO_2_ at 37°C and split at a ratio of 1:10 when reaching confluency, twice per week. The generation of HeLa ADAR1-deficient cells (HeLa 1KO) has been described previously ^37^. HeLa ADAR1/2-deficient cells (HeLa DKO) were generated using the same CRISPR/Cas9n strategy targeting ADAR2 with a mix of six guide RNA sequences (hADAR2_A^−^: GCAGCACTGATGTGAAGGAA; hADAR2_A^+^: TCTGGACAACGTGTCCCCCA; hADAR2_B^−^: GGGCCTGGCGAGGGCTCTCA; hADAR2_B^+^: AATGGGGGTGGTGGTGGCCC; hADAR2_E^−^: CTCCGAGAGCGGGGAGAGCC; hADAR2_E^+^: CAAGAGCTTCGTCATGTCTG). Single clones were selected and ADAR2 knockout was verified by western blot analysis.

### Western blot analysis and antibodies

Cells were seeded into 6 well plates and cell lysates were generated when cells reached confluency. Cells were lysed by incubating with 100 ul RIPA buffer (50 mM Tris, pH 8.0; 150 mM NaCl; 1% NP-40; 0.5% Na-deoxycholate; 0.1% SDS) supplemented with 1% protease inhibitor cocktail (Sigma-Aldrich, #P8340-5ML) on ice for 15 min. Nuclei were pelleted by centrifugation at 20,000 g, 4 °C, 15 min. Supernatants were collected, and protein content was quantified by Bradford protein assay (Bio-Rad, #5000006) using a bovine serum albumin (BSA) standard (Sigma-Aldrich, #A3803). Colorimetric measurements were obtained on a Multiskan FC Microplate Photometer (Thermo Fisher).

Samples were mixed with equal volumes of 2x Urea sample buffer (200 mM Tris-HCl, pH 6.8; 8 M Urea; 5% SDS; 0.1 mM EDTA; 0.04% bromophenol blue; 1.5% DTT) and stored at −20 °C. For immunoblotting of ADAR2, cells were lysed in 300 μl Denaturing lysis buffer (62.5 mM Tris, pH 6.8; 2% SDS; 10% glycerol; 6 M urea; 5% beta-mercaptoethanol; 0.04% bromophenol blue). 20 μg total protein per sample was subjected to SDS-PAGE on 8% polyacrylamide gels using 1× Tris/Glycine/SDS buffer (Bio-Rad, #1610772) at 100 V for 1.5 h. Proteins were transferred to PVDF membranes by wet transfer at 400 mA for 2 h on ice using 1× Tris/Glycine (Bio-Rad, #1610771) with 20% methanol. Membranes were blocked with 5% BSA (Sigma-Aldrich, #A3803) in 1×TBS (Bio-Rad, #1706435) for 1 h at room temperature. Membranes were incubated with primary antibodies in 2.5% BSA-supplemented 1× TBST at 4°C overnight. After washing 3 times with 1× TBST, membranes were incubated with HRP-conjugated antibodies in 1× TBST and at room temperature for 1 h. After washing 3 times with 1× TBST, membranes were incubated with chemiluminescence reagent (Thermo Fisher, #34580), and images were developed using a Chemidoc system (Bio-Rad). Images were processed with Image Lab software (Bio-Rad, v6.0) and Photoshop (Adobe, v26.0).

### Antibodies

Primary antibodies were rabbit anti-ADAR1 (clone D7E2M, Cell Signaling, #14175) at 1:1,000; mouse anti-ADAR2 (clone 1.3.1, Millipore Sigma, #MABE889) at 1:1,000; mouse anti-beta-actin−horseradish peroxidase (clone AC-15, Sigma-Aldrich, #A3854) at 1:20,000. Secondary antibodies were horseradish peroxidase-conjugated goat anti-rabbit IgG H&L (Jackson # 111-035-114) at 1:25,000 and horseradish peroxidase-conjugated goat anti-mouse IgG H&L (Abcam #ab205719) at 1:25,000.

### Isolation of ferret PBMCs

Animals for this study were housed in the animal facility of the Paul-Ehrlich-Institute. All procedures were in accordance with German Federal and State regulations. An animal procedures protocol authorized by the district government in Darmstadt, Hesse was active at the time experiments were performed. Ferret PBMCs were isolated from 2 ml blood drawn from the anterior vena cava from naïve male ferrets using Lithium-Heparin coated vacuettes to prevent clotting. Whole blood was centrifuged for 10 min at 800 × g at 4 °C, after which blood plasma was removed and stored at −20 °C. The remaining cellular constituents were diluted in PBS to a final volume of 4 ml and layered on top of 5 ml Histopaque-1077 (Sigma-Aldrich, #10771). After centrifugation for 30 min at 800 × g at 4 °C with acceleration and break adjusted to lowest settings, the opaque, PBMC containing phase was collected using a Pasteur pipette. Cells were washed three times in PBS to remove residual Histopaque-1077 by centrifugation for 5 min at 500 × g at 4 °C. After washing, the cells were resuspended in RPMI-1640 medium supplemented with 10% FBS, 10,000 U/l Penicillin, 10 mg/l Streptomycin, 200 mM L-Glutamine and 10 μg/ml Phytohemagglutinin-M (Sigma-Aldrich, # 11082132001) for immune cell stimulation. Cells were incubated at 37 °C, 5% atmospheric CO_2_, and 95% humidity at a cell density of 5-10×10^6^ cells/ml for 72 h.

### RNA extraction and RNA-sequencing library preparation

RNA from HeLa WT, 1KO, and DKO cells, and from ferret PBMCs was isolated using Trizol (Thermo Fisher Invitrogen, #15596026) as previously described. ^37^. Isolated total RNA was quantified using a Nanodrop spectrophotometer ND2000 (Thermo Fisher) and 5 μg RNA samples were subjected to two rounds of DNase I treatment and re-purification with phenol/chloroform/isoamyl alcohol (omitted for ferret PBMC RNA due to low RNA yields). For this, the RNA reaction in a volume of 50 μl was mixed with an equal volume of phenol/chloroform/isoamylalcohol (25:24:1, Thermo Fisher Invitrogen, #15593031) by vortexing. Aqueous and organic phases were separated by centrifugation at at max. speed and 4 °C for 1 min. The upper aqueous phase was collected and mixed with 1/9 vol. of 3M sodium acetate solution and 2.8 vol. of ethanol abs. by vortexing vigorously. After incubation for 20 min at room temperature, samples were centrifuged at 21,000 × g and 4 °C for 30 min. Supernatants were removed, pellets were washed once with 250 μl of 70% ethanol, and centrifuged at 21,000 × g and 4 °C for 2 min. Ethanol was removed completely and air-dried pellets were resuspended in 20-40 μl nuclease-free water. Samples were stored at −80 °C until further processing.

### RNA-sequencing

Illumina TrueSeq total stranded RNA libraries (Illumina) were either prepared by Azenta (Leipzig, Germany; two replicates of each HeLa cell line and ferret PBMCs), or in-house at the Paul-Ehrlich-Institute (two replicates of each HeLa cell line) using a NNSR priming-based, strand-specific library preparation protocol as described earlier ^78^. For this, total RNA with high integrity (RIN ≥10) assessed using a Fragment Analyzer (Agilent) and depleted of ribosomal RNA using the QIAseq FastSelect-RNA HMR kit (Quiagen, # 334386) was used. A full list of datasets used in this study is provided in **Supplementary Table 1**. RNA sequencing was performed by Azenta using Illumina NovaSeq 6000 (Illumina) performing 150 bp paired-end sequencing with a target read depth of 50-100 million reads per sample.

### Deposited RNA-sequencing datasets from other studies

For analysis of editing in mouse samples, we retrieved FASTQ datasets from the sequence read archive (SRA). Accession numbers for WT samples were SRR5223117, SRR5223118, and SRR5223119 from Gene Expression Omnibus (GEO) series GSE94387 ^51^; accession numbers for DKO samples were SRR9203380, SRR9203381, and SRR9203382 from GSE132214 ^52^. A full list of datasets used in this study is provided in **Supplementary Table 1**. Data were downloaded from SRA using SRATOOLKIT fasterqdump (v. 3.0.1). For analysis of editing in HeLa cells, we used samples SRR7239121 (HeLa WT) and SRR7239129 (HeLa 1KO) from GSE115127 ^37^. RNA-sequencing datasets from the Genotype-Tissue Expression Project (GTEx) were accessed through Mayo Clinic’s Bioinformatics Core agreement with the National Institutes of Health database of Genotypes and Phenotypes (dbGaP). Analyzed datasets were SRR2167030, SRR2167642, and SRR2170408 from BioProject accession number PRJNA75899 / dbGaP accession phs000424.

### Reference data for RNA-sequencing analysis

Mouse RNA-sequencing data was aligned to the mm10 reference genome (gencode.v25 / GCF_000001635.26). FASTA reference GRCm38.p6.genome.fa.gz and genome annotation gencode.vM25.annotation.gtf.gz were retrieved from https://ftp.ebi.ac.uk/pub/databases/gencode/Gencode_mouse/release_M25/.

RepeatMasker output mm10.fa.out.gz was retrieved from https://hgdownload.cse.ucsc.edu/goldenPath/mm10/bigZips/.

Human RNA-sequencing data was aligned to the hg38 reference genome (gencode.v42 / GCA_000001405.15). FASTA reference GCA_000001405.15_GRCh38_no_alt_analysis_set.fna.gz was retrieved from Indexof/genomes/all/GCA/000/001/405/GCA_000001405.15_GRCh38/seqs_for_alignment_pipelines.ucsc_ids (nih.gov). Annotation file gencode.v42.primary_assembly.annotation.gtf.gz was retrieved from https://ftp.ebi.ac.uk/pub/databases/gencode/Gencode_human/release_42/. RepeatMasker output hg38.fa.out.gz was retrieved from https://hgdownload.cse.ucsc.edu/goldenpath/hg38/bigZips/.

Ferret RNA-sequencing data was aligned to the MusPutFur1.0 reference genome (GCF_000215625.1). FASTA reference GCF_000215625.1_MusPutFur1.0_genomic.fna.gz, genome annotation GCF_000215625.1_MusPutFur1.0_genomic.gtf.gz, and RepeatMasker output GCF_000215625.1_MusPutFur1.0_rm.out.gz were retrieved from https://ftp.ncbi.nlm.nih.gov/genomes/all/GCF/000/215/625/GCF_000215625.1_MusPutFur1.0/. For downstream analysis with Bedtools, genome annotation files were split into gene and exon/UTR annotation files. An intron annotation file was generated by strand-specific subtraction of exon/UTR regions from gene regions using BEDTOOLS subtract (v. 2.30.0) ^79^. RepeatMasker output files were converted into BED format using BEDOPS rmsk2bed (v. 2.4.41) ^80^.

### RNA-sequencing alignment pipeline

All processes were performed in the mforge computational cluster of Mayo Clinic, where individual programs are available as modules. FASTQ files separated by read group were quality-trimmed and filtered short reads using fastp (v. 0.23.1) ^81^ using parameters [-M 30 −3 −5 -n 0 −l 35 -q 30 -u 15 -w 16 -f 1 -F 1]. Processed FASTQ files were then aligned by STAR (v. 2.7.9a) ^82^using parameters [--runThreadN 30 -- outSAMmapqUnique 60--outSAMtype BAM SortedByCoordinate--twopassMode Basic -- outFilterType Normal--outFilterMultimapNmax 10--alignSJoverhangMin 5 -- alignSJDBoverhangMin 3--outFilterMismatchNmax 10--outFilterMismatchNoverReadLmax 1--outFilterScoreMinOverLread 0--outFilterMatchNminOverLread 0--outFilterMatchNmin 0--alignIntronMin 21--alignIntronMax 0--alignMatesGapMax 0]. Next, a read group identifier was added with GATK AddOrReplaceReadGroups (v. 4.3.0.0) ^83^ using parameters--RGID $sampleID--RGLB $sampleID--RGPL illumina--RGPU 0--RGSM $sampleID -- SORT_ORDER coordinate]. Optical duplicates were marked and removed with PICARD MarkDuplicates (v. 2.27.5) ^84^ using parameters [CREATE_INDEX=true REMOVE_DUPLICATES=true VALIDATION_STRINGENCY=SILENT]. Reads spanning exon/exon junctions were split using GATK SplitNCigarReads (v. 4.3.0.0) ^83^ using default settings, and re-aligned with GATK LeftAlignIndels (v. 4.3.0.0) ^83^ using parameter [--disable-read-filter WellformedReadFilter]. At each step, bai index files were created using SAMTOOLS index (v. 1.16) ^85^, and bam file statistics were calculated with SAMTOOLS stats (v. 1.16) ^85^. To combine BAM alignments from biological replicates, we employed a pipeline of BAMTOOLS merge, sort, and index (v. 2.5.2) ^86^ using default settings.

### TSniffer analysis

The *TSniffer* package can be found at https://github.com/maiher/tsniffer_1.0. A yml file to create a conda environment with all dependencies to run *TSniffer* is provided. [The *TSniffer* code will be made freely available for non-commercial use upon final publication of the manuscript. During the peer review process, reviewers will be provided access to the package and instructions to install the *TSniffer* conda environment. A test dataset will be provided for analysis.] *TSniffer deNovo* requires a BAM alignment file and the corresponding reference genome as input. It generates read count tables and calculates the occurrence of transition mutation types (TsTypes): A>G (AG), C>T (CT), G>A (GA), and T>C (TC), and transversion (or non-transition) mutation types (A>C, A>T, C>A, C>G, G>C, G>T, T>A, T>G) dependent on each reference base (A,C, G, T) separately at each position. Using a window of user-defined size (default:-wSize 100), it then performs a sliding window analysis to define regions of significant enrichment of one type of query Ts over the other types utilizing Fisher’s exact tests. Thereby, the contingency matrix contains the accumulated counts of reads exhibiting query transitions (q-Ts), non-query transitions (nq-Ts), query non-transitions (q-nTs), and non-query non-transitions (nq-nTs) across the window. A user-defined minimum of coverage can be applied for a site to be included (default:-minCov 4). To assist computational needs, chromosomes can be split into overlapping chunks of defined sizes to be analyzed consecutively (default:-chunkSize 50,000,000). Analysis can be restricted to specific chromosomes (-chr chrxyz) or primary assemblies (default:-primary). The analyzed TsTypes can be specified (-tsType AG CT GA TC all, default: AG TC). We used *TSniffer deNovo* with following parameters [-wSize 100-minCov 4-chunkSize 5,000,000-primary (omitted for ferret data)]. Each TsType was analyzed in a separate run to allow easier downstream analysis. TsRegions were saved in GFF files, separated by chromosome/contig. For easier downstream analysis, individual GFF files for each TsType were concatenated. The output file contains Ts types (type) mean and exact relative transition frequencies (rtf_mean / rtf_exact), mean and exact *p*-values from Fisher’s exact test (p_mean / p_exact), number of identified Ts sites (TsSiteCount), and exact coordinates of Ts sites (TsPos). Mean values are calculated from individual windows merged into a TsRegion, whereas exact values are calculated by re-analyzing the final significant TsRegion. For downstream analyses, we utilized rtf_exact values and TsSiteCounts. To achieve highest sensitivity, *TSniffer deNovo* analysis was performed on combined BAM alignments of biological replicate samples.

*TSniffer Regio* analysis requires a BAM alignment file, the corresponding reference genome, and a GFF file with regions to be analyzed (e.g. output from *TSniffer deNovo*) as input. Default settings were used for our analyses described here [-wSize 100-minCov 8]. The output file was a single GFF per sample with all analyzed TsRegions. The program reports an error message in case of low coverage of specific TsRegions in the analyzed dataset: not analyzed: no position has min. coverage.

### TsRegion filtering

To minimize false-positive TsRegions arising from single nucleotide variants or sequencing/alignment artifacts, we employed the following pipeline: First, *TSniffer deNovo* TsRegions of the same type within 100 nt distance were merged into single TsRegions using BEDTOOLS merge [-d 100] ^79^. Exact RTF values and TsSiteCounts were determined by *TSniffer Regio* analysis. For analyses where ADAR-deficient datasets were available (mouse, HeLa), *TSniffer Regio* analysis was also performed on knockout (KO) datasets. Next, TsRegions found in both WT and KO datasets were removed from downstream analysis if they had 50% overlap in both datasets using BEDTOOLS subtract [-f 0.5] ^79^. This step was omitted, if no KO dataset was available (GTEx brain and ferret samples). The resulting set of TsRegions was further filtered for TsSiteCount ≥ 5, and all TsRegions with < 5 TsSites were discarded. For mouse and HeLa TsRegions, a confidence indicator (CI) was calculated based on the reduction of rtf_exact between the WT and KO datasets as described in equation 1:

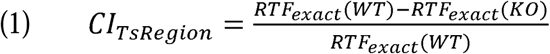

TsRegions were further filtered for different CI ranges, and a final set was determined that contained all TsRegions with CI ≥ 0.85 and TsSiteCount ≥ 5).

### Association of TsRegions with genetic elements

We determined the overlap of TsRegions of interest with different annotated genetic elements (genes, exons/UTRs, introns, repeat elements) using BEDTOOLS intersect [-wa-wb] ^79^. Downstream analysis was performed using Microsoft Excel 365 (v. 2407). Target TsRegions (AG, TC) and non-target TsRegions (CT, GA) were analyzed separately, where indicated. For analysis of overlap of TsRegions with repeat elements and exon/UTR/introns, we simply counted the number of TsRegions intersecting with specific types of genetic elements.

To calculate the overall editing scores of specific transcripts, we summed rtf_exact values of all target TsRegions overlapping with the annotated gene and, in case of available KO data, subtracted the residual background detected in these datasets, according to equation 2:

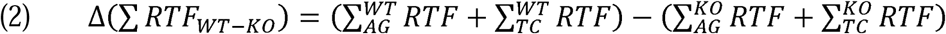

Where KO data was not available, we omitted the corrective term in equation 2. Summed TsSiteCounts were calculated accordingly.

### *LoDEI* analysis and comparison with *TSniffer*

*LoDEI* ^50^ was installed as instructed at Github (https://github.com/rna-editing1/lodei). The HeLa WT_merged and DKO_merged BAM files were analyzed against each other using –min_coverage 4 and -w 50 parameters to achieve highest possible similarity to the parameters used for *TSniffer* analyses. The resulting txt file of q-filtered significant AG windows (no other mutation type resulted in significant windows) was converted into gff format. To allow for better direct comparison with TsRegions, adjacent windows were merged into *LoDEI* regions using BEDTOOLS merge (-d 1). The overlap between *LoDEI* regions and TsRegions was analyzed by BEDTOOLS intersect. For calculation of overall editing of transcripts, window editing indices (wEI) of all windows associated with a gene (*LoDEI* includes this information directly in the output file) were summarized as described for RTF above (equation 2).

### Data and figure graphing

Diagrams of analyzed data were generated with Microsoft Excel 365 or with Graphpad Prism (v. 9.5.1). Figures were generated with Adobe Illustrator (v. 27.7).

## Data availability

Newly generated sequencing data will be deposited in GEO and SRA under BioProject #TBD. Analyzed data will be found under accession number #TBD and raw FASTQ files will be accessible under accession numbers #TBD.

## Author Contribution

MH – conceptualization, software development, algorithm testing and verification, data analysis, manuscript writing; YK – generation of cell lines, RNA extraction; FMA – data analysis, manuscript editing; SP – western blot, data analysis, manuscript editing; OS – ferret experiments, manuscript editing; FGMA – ferret PBMC purification, RNA extraction, manuscript editing; BT – data analysis, manuscript editing; CM – RNA-sequencing, manuscript editing; CKP – conceptualization, data analysis, manuscript writing, funding acquisition.

## Acknowledgements

The authors would like to thank Charles “Chuck” Samuel, Carl Walkley, Roberto Cattaneo, Bevan Sawatsky, and Katayoun Ayasoufi for their insightful comments and discussions during preparation of the manuscript. They also would like to thank Zhifu Sun and Mayo Clinic’s Bioinformatics Core (Daniel O’Brien) for granting access to GTEx datasets analyzed in this study.

---

**Extended Data Fig. 1:**
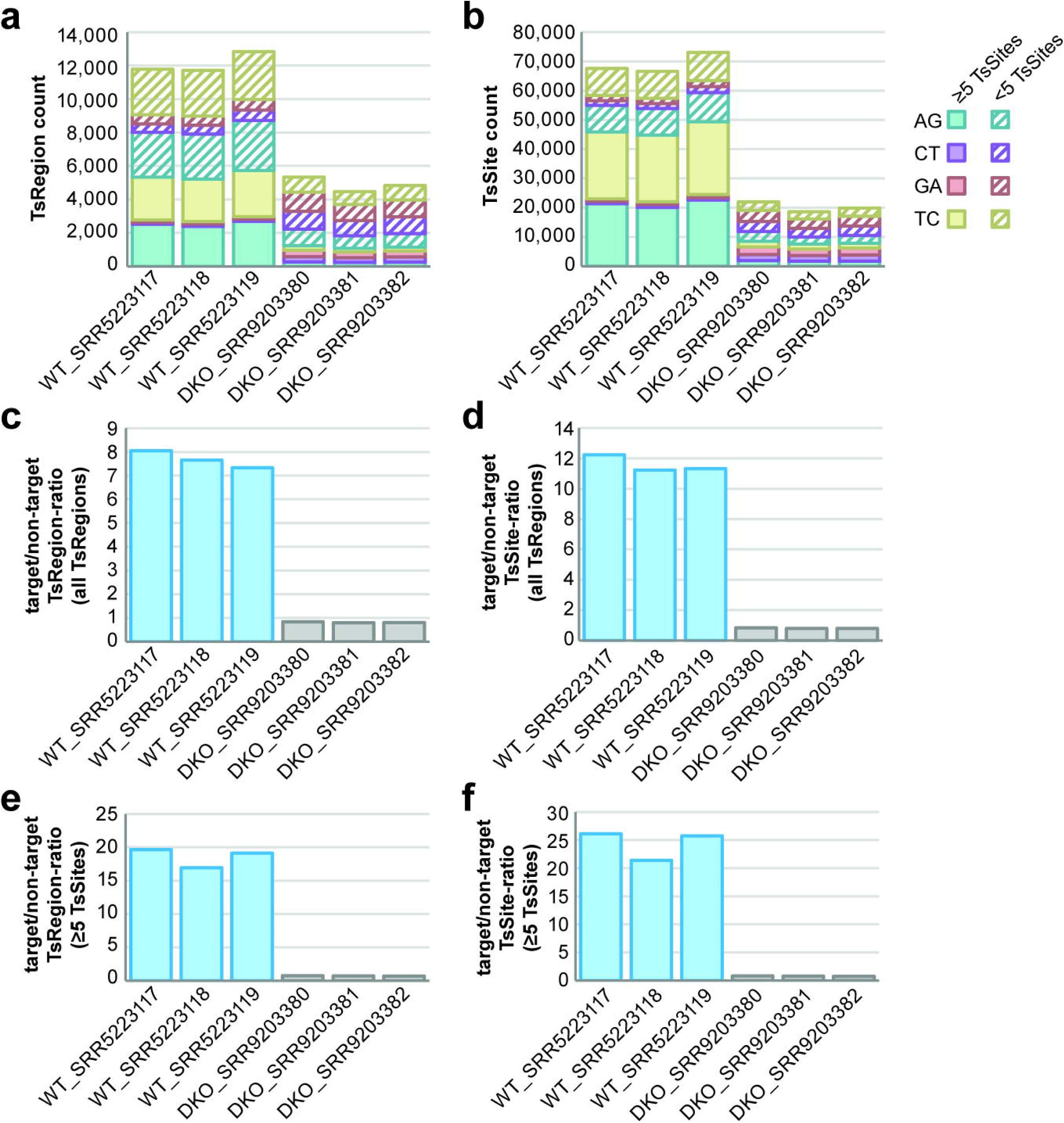
*TSniffer de novo* analysis of individual WT and DKO mouse brain datasets. (**a**) TsRegion count and (**b**) TsSite count of *de novo*-identified TsRegions in three individual BAM alignments of wild type (WT) and ADAR1/2-deficient (DKO) mouse brains. (**c**) Ratio of target (AG + TC) to non-target (CT + GA) TsRegion counts in each dataset. (**d**) Ratio of target (AG + TC) to non-target (CT + GA) TsSite counts in each dataset. (**e**) Ratio of target (AG + TC) to non-target (CT + GA) TsRegion counts in each dataset. Only TsRegions with at least 5 TsSites are included in this calculation. (**f**) Ratio of target (AG + TC) to non-target (CT + GA) TsSite counts in each dataset. Only TsSites from TsRegions with at least 5 TsSites are included in this calculation.

**Extended Data Fig. 2:**
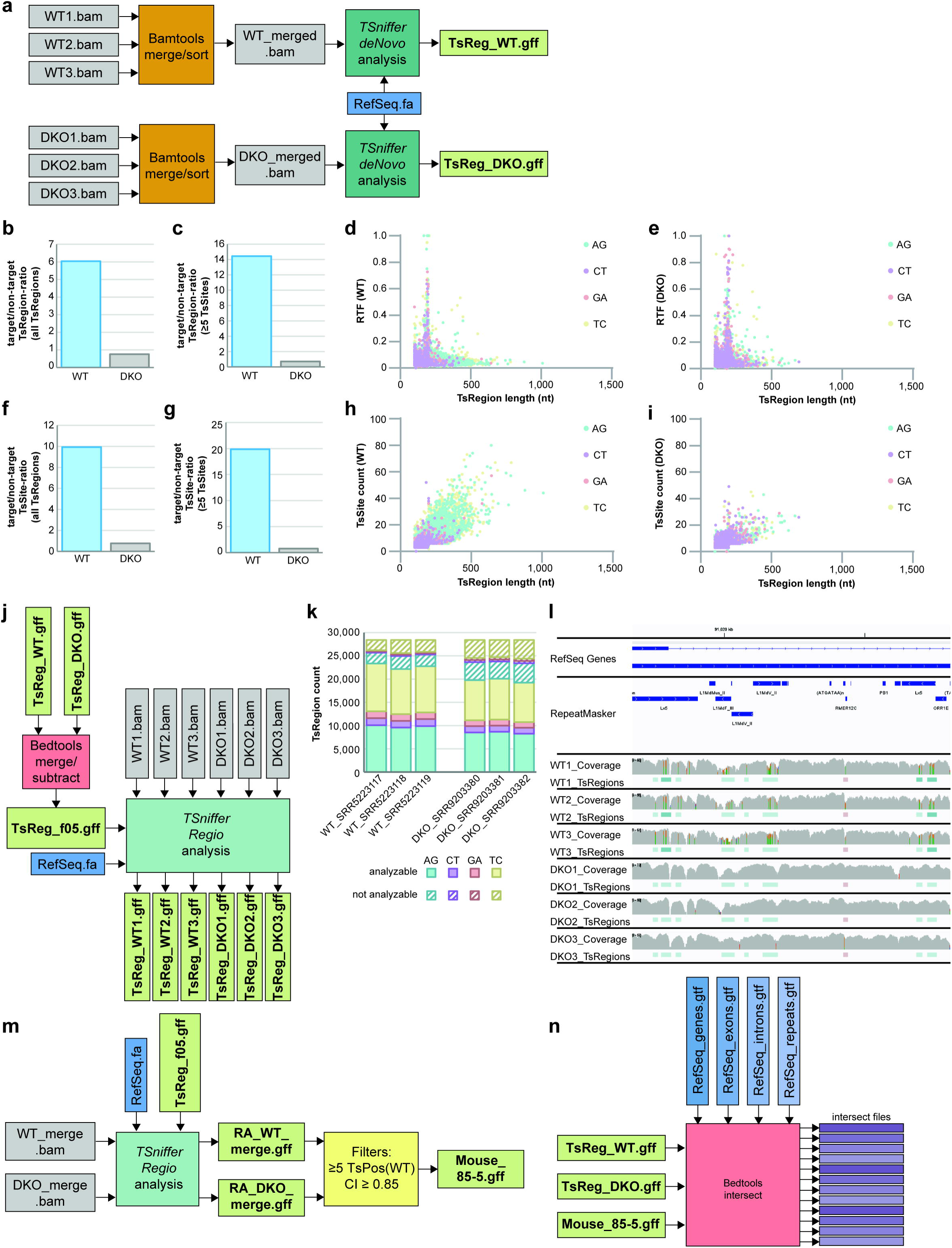
Optimized *TSniffer* pipeline and downstream analysis. (**a**) Workflow of *TSniffer deNovo* analysis on combined BAM alignments from biological replicates. (**b**) Ratio of target (AG + TC) to non-target (CT + GA) TsRegion counts in combined BAM datasets. (**c**) Ratio of target (AG + TC) to non-target (CT + GA) TsRegion counts in combined BAM datasets. Only TsRegions with at least 5 TsSites are included in this calculation. (**d**) Distribution of TsRegion length and RTF values of TsRegions in the combined WT dataset. Different TsTypes are color-coded. (**e**) Distribution of TsRegion length and RTF values of TsRegions in the combined DKO dataset. (**f**) Ratio of target (AG + TC) to non-target (CT + GA) TsSite counts in combined BAM datasets. (**g**) Ratio of target (AG + TC) to non-target (CT + GA) TsSite counts in combined BAM datasets. Only TsSites of TsRegions with at least 5 TsSites are included in this calculation. (**h**) Distribution of TsRegion length and TsSite counts of TsRegions in the combined WT dataset. Different TsTypes are color-coded. (**i**) Distribution of TsRegion length and TsSite counts of TsRegions in the combined DKO dataset. (**j**) Workflow for removal of TsRegions shared between WT and DKO samples (TsReg_f05) and downstream *TSniffer Regio* analysis of individual BAM alignments with this set of TsRegions. (**k**) TsRegion count of *TSniffer Regio* analysis of individual BAM alignments with the TsReg_f05 set. Hashed bars indicate proportion of TsRegions returning no analyzable value due to low coverage in the alignment. (**l**) Screenshot of an alignment of a transcript region in IGV. Coverage plots (allele frequency threshold set to 0.05) and *TSniffer Regio* analysis GFF files were loaded. Darker colors of TsRegions indicate higher RTF values. (**m**) Workflow for establishing a highly representative and specific subset of TsRegions (Mouse_85-5). For details see methods. (**n**) Workflow for downstream BEDTOOLS analysis of identified TsRegions.

**Extended Data Fig. 3:**
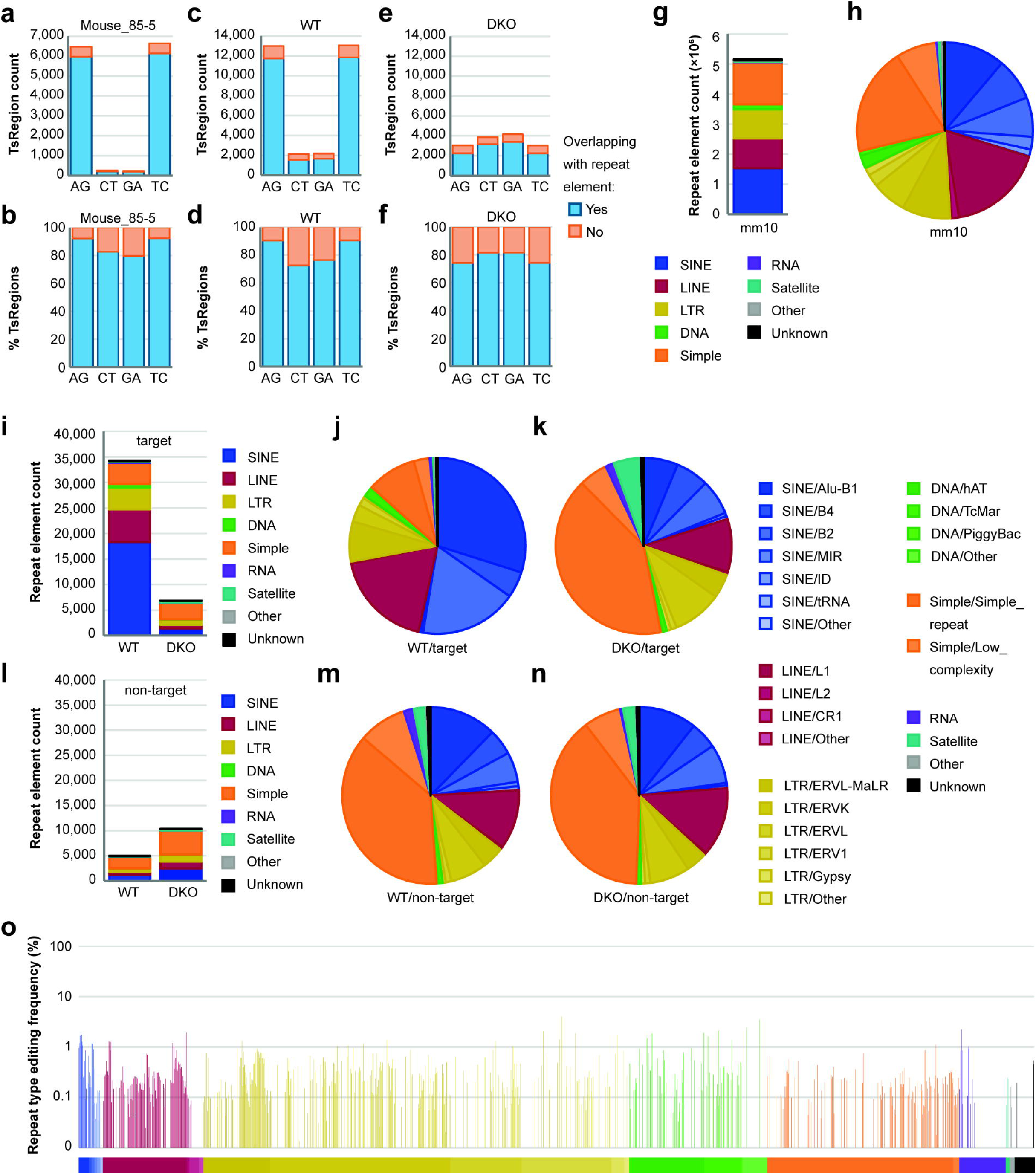
ADAR editing in murine repeat elements. (**a**) Mouse_85-5 TsRegion count on intersection with annotated repeat elements. (**b**) Relative frequencies of Mouse_85-5 TsRegions overlapping with repeat elements. (**c**) Mouse WT *de novo* TsRegion count on intersection with annotated repeat elements. (**d**) Relative frequencies of Mouse WT *de novo* TsRegions overlapping with repeat elements. (**e**) Mouse DKO *de novo* TsRegion count on intersection with annotated repeat elements. (**f**) Relative frequencies of Mouse DKO *de novo* TsRegions overlapping with repeat elements. (**g**) Total repeat element count and (**h**) relative occurrence of repeat element subtypes in the mouse reference genome mm10. (**i**) Total counts of repeat element families overlapping with target TsRegions (AG + TC) in WT and DKO *de novo* datasets. (**j**) Relative frequencies of mouse repeat subtypes overlapping with target TsRegions in the WT *de novo* dataset. (**k**) Relative frequencies of mouse repeat subtypes overlapping with target TsRegions in the DKO *de novo* dataset. (**l**) Total counts of repeat element families overlapping with non-target TsRegions (CT + GA) in WT and DKO *de novo* datasets. (**m**) Relative frequencies of mouse repeat subtypes overlapping with non-target TsRegions in the WT *de novo* dataset. (**n**) Relative frequencies of mouse repeat subtypes overlapping with non-target TsRegions in the DKO *de novo* dataset. (**o**) Frequencies of TsRegion-harboring repeat elements by specific repeat type. Color coding as in **i** – **n**.

**Extended Data Fig. 4:**
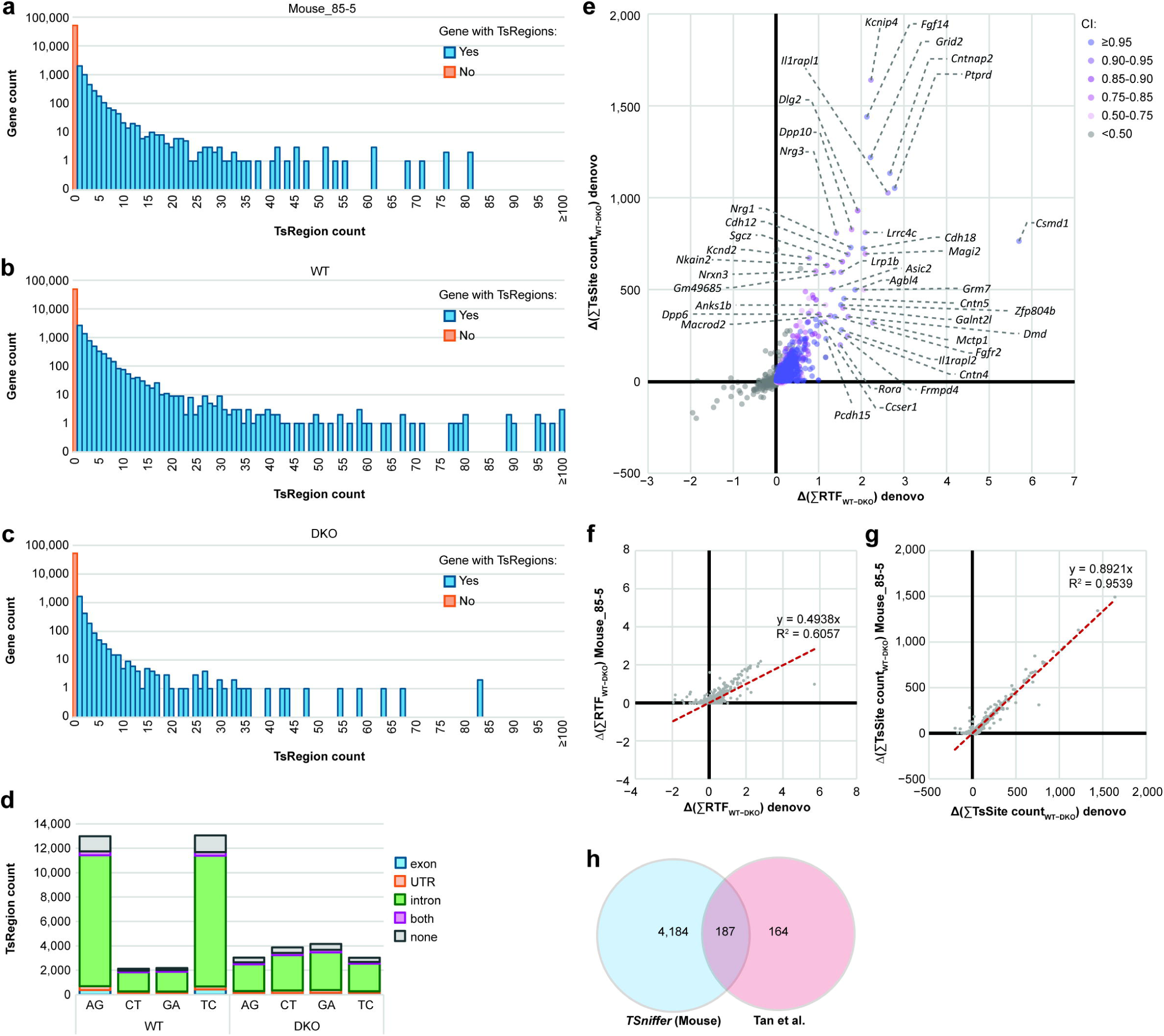
Consistency between *TSniffer de novo* and Mouse_85-5 subset analyses. (**a**) Counts of genes harboring specific numbers of TsRegions of the Mouse_85-5 set. (**b**) Counts of genes harboring specific numbers of TsRegions of the Mouse WT *de novo* set. (**c**) Counts of genes harboring specific numbers of TsRegions of the Mouse DKO *de novo* set. (**d**) Distribution of WT and DKO *de novo* TsRegions across transcript elements. (**e**) Quantification of ADAR editing in mouse transcripts. Relative transition frequency (RTF) values and TsSite counts of all Mouse WT *de novo* TsRegions intersecting with individual genes were summed up and values of respective TsRegions in DKO were subtracted. Transcripts are color-coded by total number of calculated TsSites. Most highly edited transcripts are indicated. (**f**) Correlation of gene-specific RTF values derived from the Mouse_85-5 set and from the subtractive *de novo* analysis. (**g**) Correlation of gene-specific TsSite counts derived from the Mouse_85-5 set and from the subtractive *de novo* analysis. (**h**) Meta-analysis of edited genes detected by *TSniffer* in mouse brain samples and by Tan et al.

**Extended Data Fig. 5:**
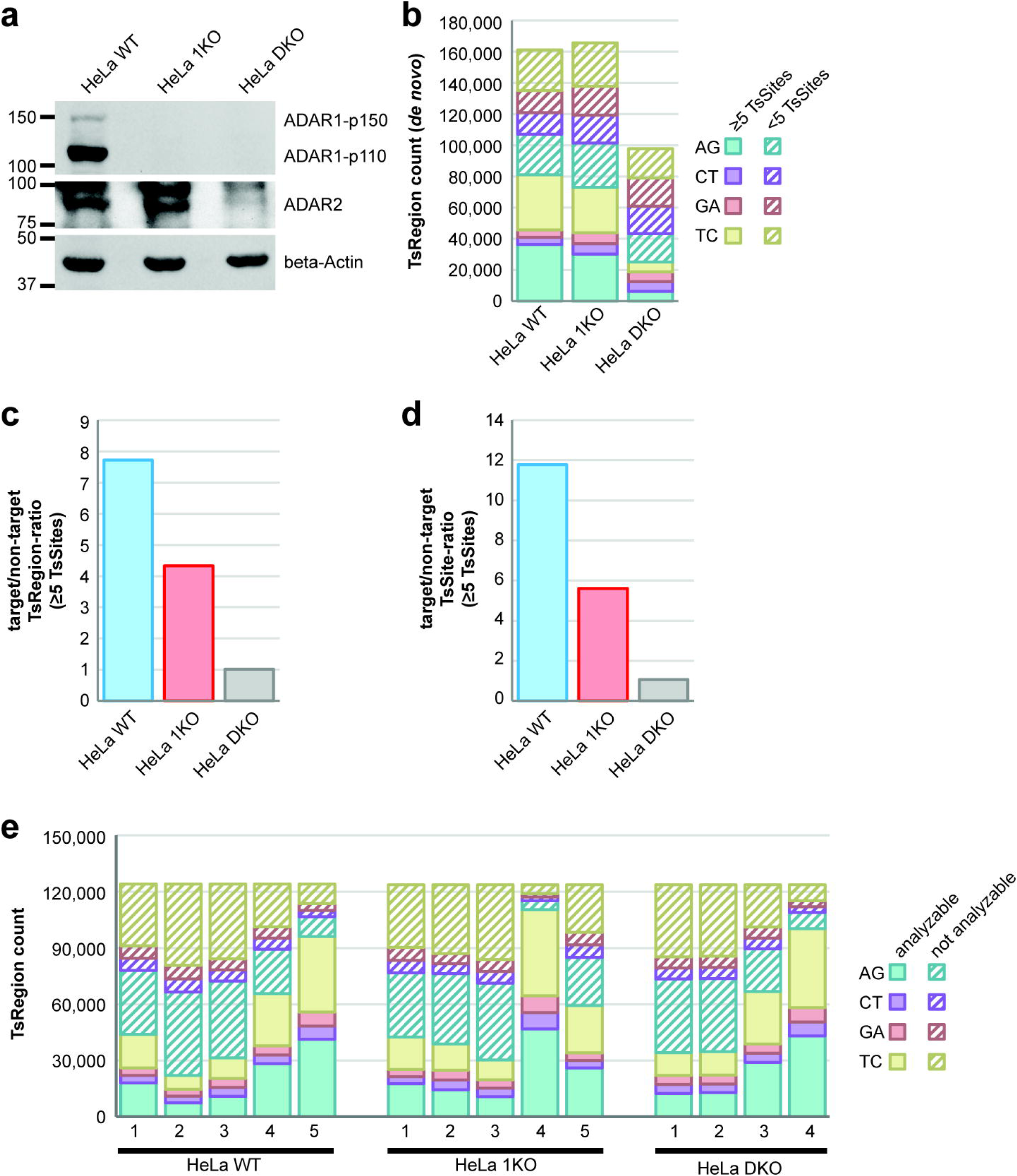
Generation of ADAR1- and ADAR1/2-deficient HeLa cell lines and initial *TSniffer* analysis. (**a**) Western blot analysis of wild type (WT), ADAR1 knockout (1KO) and ADAR1/2 knockout (DKO) HeLa cells. (**b**) TsRegion count of *TSniffer de novo* analyses of combined datasets of BAM alignments of biological replicates of HeLa WT (n=5), HeLa 1KO (n=5), and HeLa DKO (n=4). (**c**) Ratio of target (AG + TC) to non-target (CT + GA) TsRegion counts in combined BAM datasets. Only TsRegions with at least 5 TsSites are included in this calculation. (**d**) Ratio of target (AG + TC) to non-target (CT + GA) TsSite counts in combined BAM datasets. Only TsSites of TsRegions with at least 5 TsSites are included in this calculation. (**e**) *TSniffer* region anlysis of individual BAM alignments derived from HeLa WT (n=5), HeLa 1KO (n=5), and HeLa DKO (n=4) using the HeLa WT-specific set of TsRegions. Hashed bars indicate TsRegions returning non-analyzable values in each dataset due to low coverage.

**Extended Data Fig. 6:**
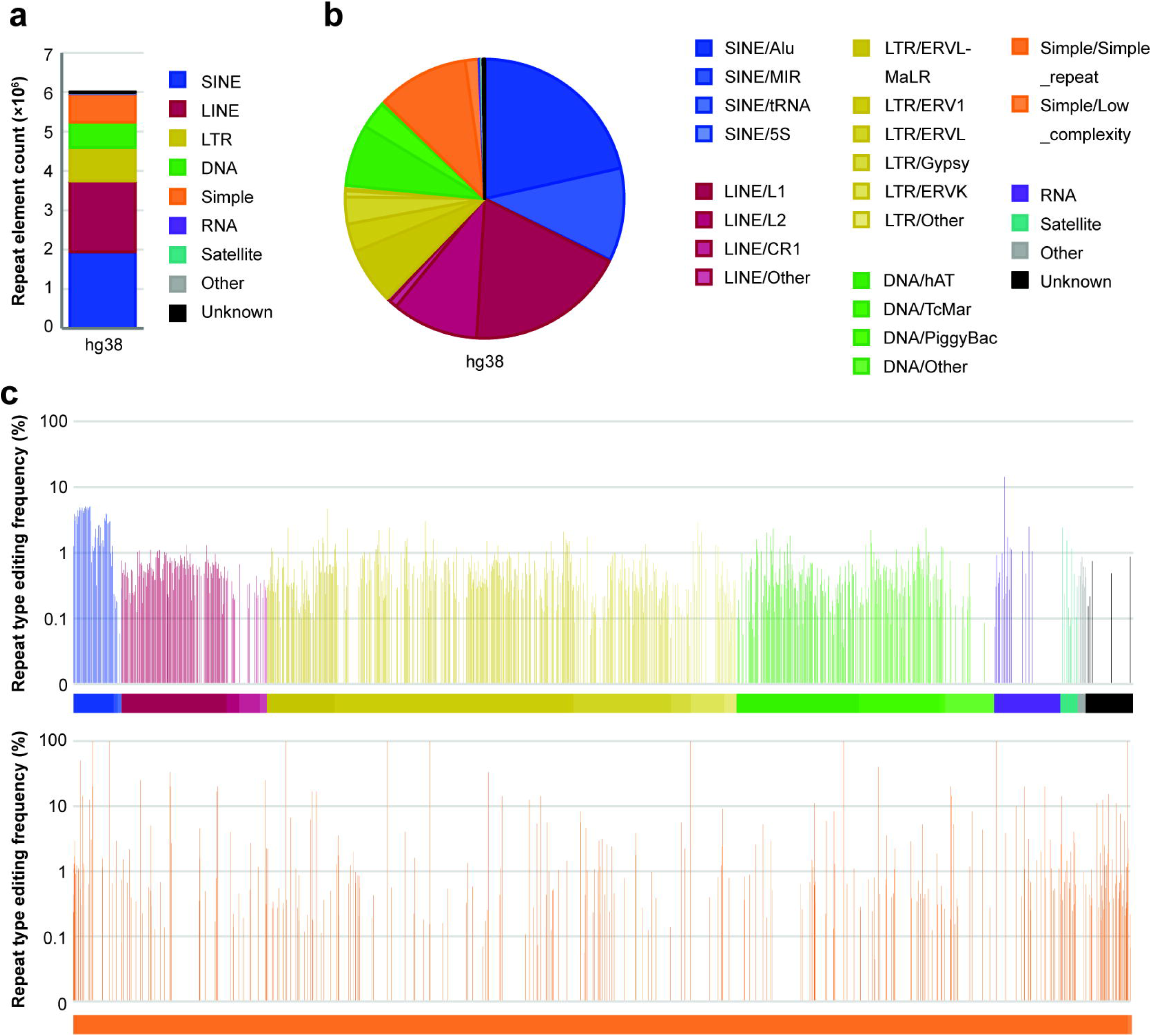
Human repeat elements and frequency of ADAR editing within these elements. (**a**) Total repeat element count and (**b**) relative occurrence of repeat element subtypes in the human reference genome hg38. (**c**) Frequencies of TsRegion-harboring repeat elements by specific repeat type. Color coding as in **a** – **b**. Due to the high number of annotated simple repeats in hg38, these are shown in a separate diagram (bottom).

**Extended Data Fig. 7:**
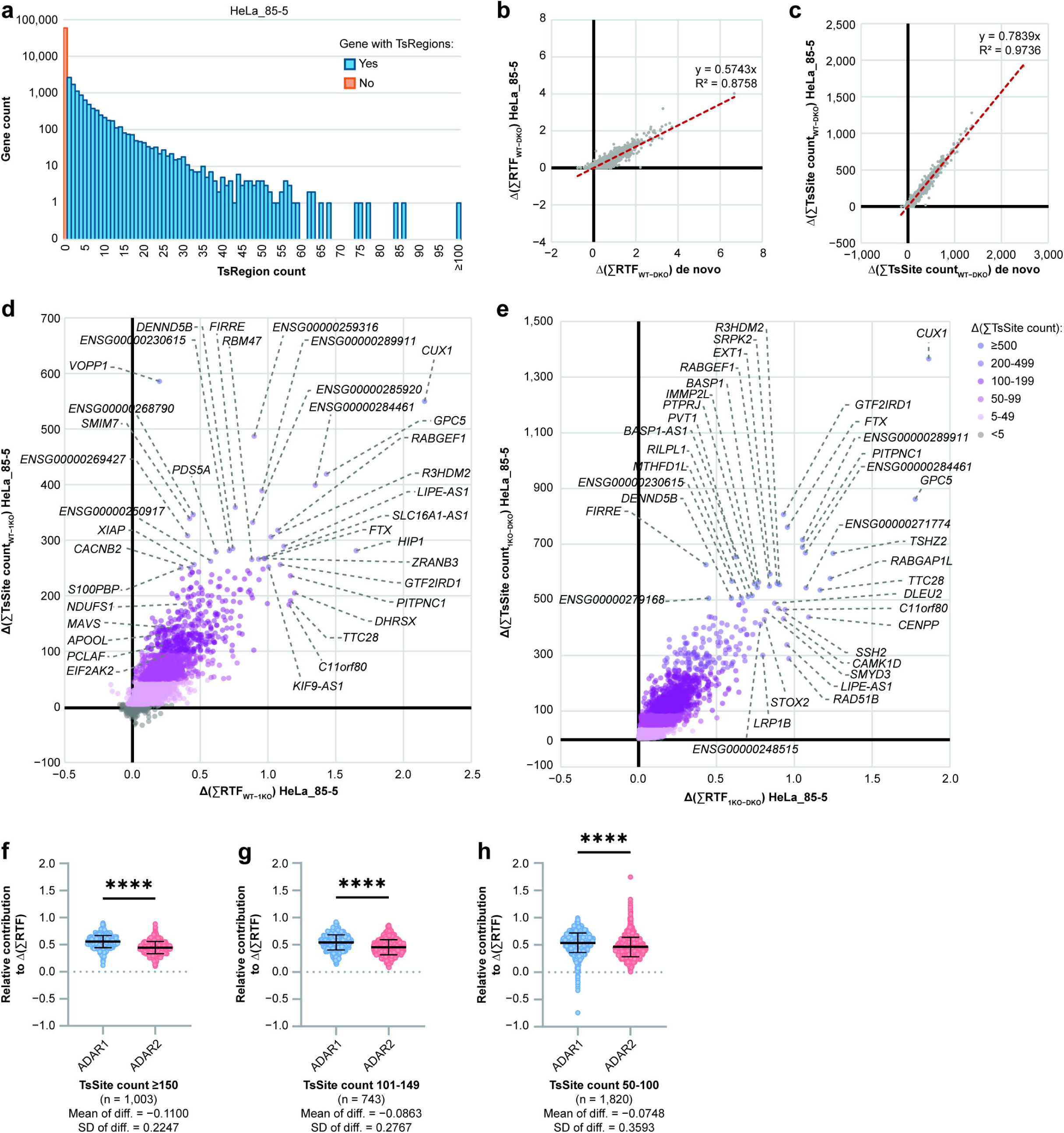
Differential editing of ADAR1 and ADAR2 in human transcripts. (**a**) Counts of genes harboring specific numbers of TsRegions of the HeLa_85-5 set. (**b**) Correlation of gene-specific RTF values derived from the Mouse_85-5 set and from the subtractive *de novo* analysis. (**c**) Correlation of gene-specific TsSite counts derived from the Mouse_85-5 set and from the subtractive *de novo* analysis. (**d**) Quantification of the ADAR1-specific portion of editing in human transcripts. Relative transition frequency (RTF) values and TsSite counts of all HeLa_85-5 TsRegions intersecting with individual genes in WT were summed up and residual values of respective regions in 1KO were subtracted. Transcripts are color-coded by total number of calculated TsSites. Most highly edited transcripts are indicated. (**e**) Quantification of the ADAR2-specific portion of editing in human transcripts. Relative transition frequency (RTF) values and TsSite counts of all HeLa_85-5 TsRegions intersecting with individual genes in 1KO were summed up and residual values of respective regions in DKO were subtracted. Transcripts are color-coded by total number of calculated TsSites. Most highly edited transcripts are indicated. (**f**) Comparison of contribution of ADAR1 and ADAR2 to RTF values of genes with at least 150 TsSites (n = 1,003). (**g**) Comparison of contribution of ADAR1 and ADAR2 to RTF values of genes with 101-149 TsSites (n = 743). (**h**) Comparison of contribution of ADAR1 and ADAR2 to RTF values of genes with 50-100 TsSites. In **f** – **h**, paired two-tailed Student’s t-test was performed between groups to test for significant differences. **** *P*<0.0001.

**Extended Data Fig. 8:**
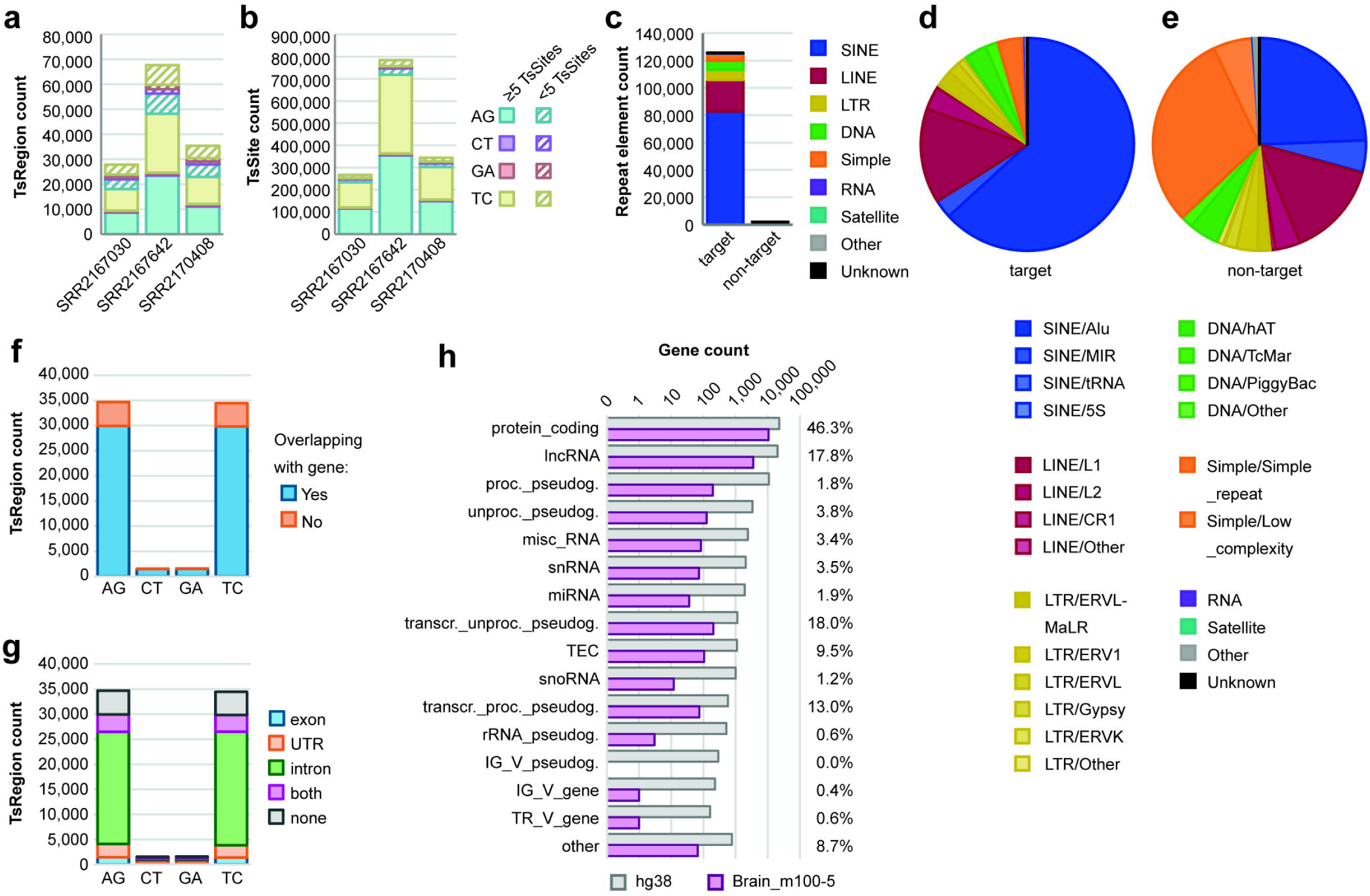
ADAR editing in GTEx datasets of primary human brain samples. (**a**) TsRegion counts and (**b**) TsSite counts of *TSniffer deNovo* analyses of three GTEx brain samples. (**c**) Number of repeat elements overlapping with TsRegions of the Brain_m100-5 set. Target TsRegions are AG and TC, non-target TsRegions are CT and GA. Colors indicate different repeat families. (**d**) Relative abundance of repeat family subtypes among target TsRegion-containing repeats. (**e**) Relative abundance of repeat family subtypes among non-target TsRegion-containing repeats. (**f**) TsRegion count by intersection with annotated genes. (**g**) Distribution of TsRegions within transcript elements. (**h**) Total number of annotated genes in the hg38 reference genome (grey), total number of genes with target TsRegions (pink), and relative frequency of TsRegion-harboring genes, per gene type.

**Extended Data Fig. 9:**
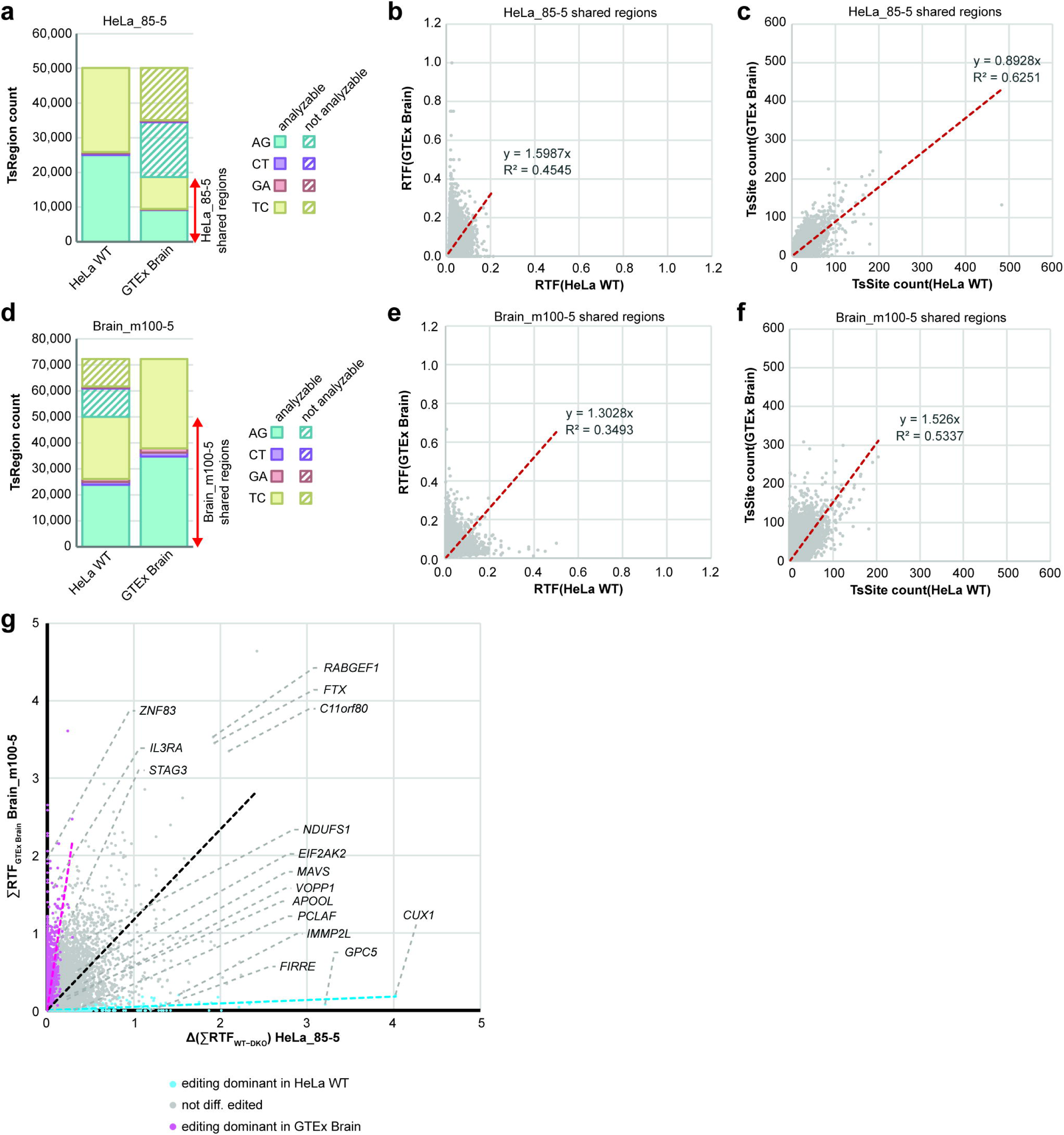
Comparison of ADAR editing in HeLa cells and primary human brain tissue. (**a**) TsRegion count of *TSniffer Regio* analysis of HeLa WT and GTEx Brain datasets using the HeLa_85-5 set of TsRegions. Hashed bars indicate TsRegions returning not analyzable values in the respective dataset due to low coverage. Red arrow indicates the subset of TsRegions used for correlation analysis in **b** and **c**. (**b**) Correlation of RTF values obtained from HeLa WT cells (x-axis) and the GTEx Brain dataset (y-axis) for HeLa_85-5 TsRegions that were expressed in both datasets. (**c**) Correlation of TsSite counts obtained from HeLa WT cells (x-axis) and the GTEx Brain dataset (y-axis) for HeLa_85-5 TsRegions that were expressed in both datasets. (**d**) TsRegion count of *TSniffer Regio* analysis of HeLa WT and GTEx Brain datasets using the Brain_m100-5 set of TsRegions. Hashed bars indicate TsRegions returning not analyzable values in the respective dataset due to low coverage. Red arrow indicates the subset of TsRegions used for correlation analysis in **e** and **f**. (**e**) Correlation of RTF values obtained from HeLa WT cells (x-axis) and the GTEx Brain dataset (y-axis) for Brain_m100-5 TsRegions that were expressed in both datasets. (**f**) Correlation of TsSite counts obtained from HeLa WT cells (x-axis) and the GTEx Brain dataset (y-axis) for Brain_m100-5 TsRegions that were expressed in both datasets. (**g**) Correlation of gene-associated total RTF values detected in HeLa WT cells using the HeLa_85-5 set (x-axis) with values from human brain samples using the Brain_m100-5 set (y-axis). Color coding if gene was dominantly edited in HeLa WT (blue; TsSite count[HeLa WT] ≥ 9 × TsSite count[Brain]) or if gene was dominantly edited in brain (pink; TsSite count[Brain] ≥ 9 × TsSite count[HeLa WT]. Dashed lines indicate linear regression for the different groups.

**Extended Data Fig. 10:**
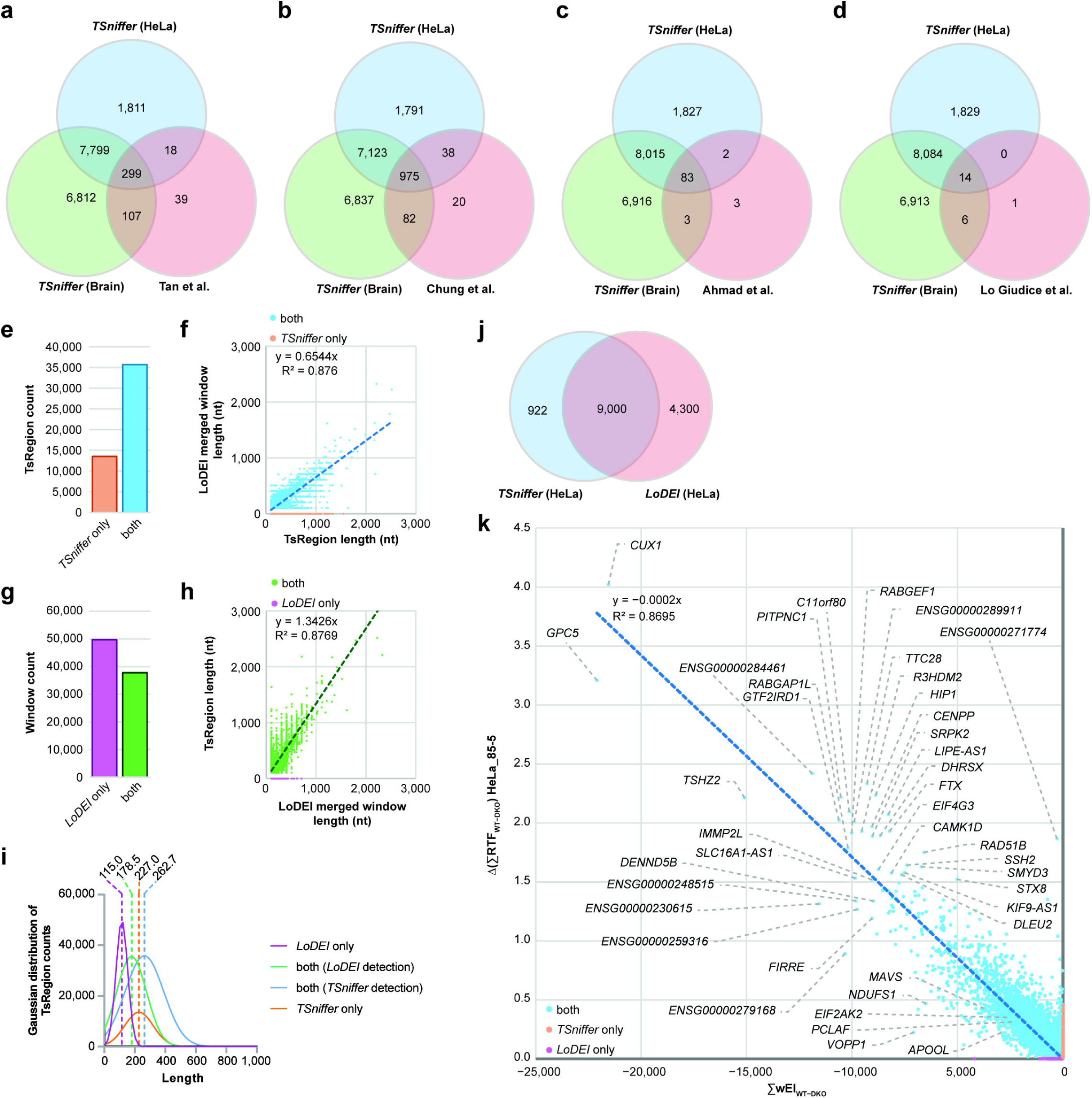
Performance comparison of *TSniffer* with other approaches to detect and quantify RNA editing. (**a**) Meta-analysis of edited genes detected by *TSniffer* in HeLa or GTEx brain samples and by Tan et al. (**b**) Meta-analysis of edited genes detected by *TSniffer* in HeLa or GTEx brain samples and by Chung et al. (**c**) Meta-analysis of edited genes detected by *TSniffer* in HeLa or GTEx brain samples and by Ahmad et al. (**d**) Meta-analysis of edited genes detected by *TSniffer* in HeLa or GTEx brain samples and by Giudice et al. (**e**) Number of HeLa_85-5 TsRegions detected by *TSniffer* only (orange) or by *TSniffer* and *LoDEI* (blue). (**f**) Comparison of the TsRegion length calculated by *TSniffer* and the cumulative length of overlapping significant windows detected by *LoDEI*. (**g**) Number of merged adjacent windows detected by *LoDEI* only (purple) or by *LoDEI* and *TSniffer* (green). (**h**) Comparison of the cumulative length of windows detected by *LoDEI* and overlapping TsRegions calculated by *TSniffer*. (**i**) Gaussian distribution of window/TsRegion length in all four groups (**f** – **h**). Dashed lines and values on top indicate the mean region length for each group. (**j**) Number of gene transcripts in HeLa cells identified by *TSniffer* (HeLa_85-5 set), or *LoDEI*, and overlap between the two analyses. (**k**) Correlation of the cumulative window editing index (wEI) for each transcript as calculated by *LoDEI* between HeLa WT and DKO samples, and the cumulative RTF values as calculated by *TSniffer* for the same datasets.

